# Comprehensive Longitudinal ctDNA Monitoring in Metastatic Cancer Patients Treated with an Individualized Neoantigen-directed Vaccine

**DOI:** 10.1101/2024.12.04.626817

**Authors:** Desiree Schenk, Matthew J. Davis, Rita Zhou, Alexis Mantilla, Madeline Galbraith, Oliver Spiro, Olivia Petrillo, Italo Faria do Valle, Andrew R. Ferguson, Karin Jooss, Ankur Dhanik

## Abstract

**Purpose:** Circulating-tumor DNA (ctDNA) is an emerging, minimally invasive diagnostic and prognostic biomarker for patients receiving a variety of cancer therapies. Comprehensive and robust longitudinal monitoring of ctDNA can provide an understanding of tumor burden, heterogeneity, and response or resistance to treatment.

**Experimental Design:** ctDNA of 28 metastatic cancer patients receiving an individualized neoantigen-directed immunotherapy was monitored longitudinally, up to two years, using a unique hybrid next generation sequencing assay targeting tumor-informed and tumor-naïve variants. Patient-specific panels were designed targeting an average of 144 variants per patient. A tumor-naïve universal panel was also designed for inclusion with patient-specific panels to monitor recurrently mutated tumor hotspots (e.g., *KRAS* and *TP53*) and genes implicated in immunotherapy resistance (*B2M, TAP1/2*).

**Results:** Analytical characterization of the assay established linearity with a mean variant allele frequency (VAF) ≥0.049%, and a variant-level limit of detection (LOD95) of 0.12%. Tumor-informed variants were detected in 26/28 patients, and *de novo* variants were observed in 25/28 patients. HLA LOH was also observed. Longitudinal ctDNA data provided key insights into patients’ responses to vaccine treatment.

**Conclusions:** The hybrid design of the ctDNA monitoring assay provides the sensitivity and specificity required for evaluating patient samples undergoing individualized therapy. It provides an improved capability to understand patient response to experimental therapies and further supports the utility of ctDNA as a cancer biomarker.

## Introduction

Cell-free DNA (cfDNA) from liquid biopsies has become an important analyte for longitudinal disease monitoring in oncology patients. Unlike traditional biopsies, liquid biopsies are easy to collect via a simple venipuncture and offer the opportunity for more frequent sampling than tissue biopsies. From a prognostic standpoint, patients with lower ctDNA tend to have lower disease burden and better clinical outcomes than those with higher ctDNA. Aside from its prognostic potential, ctDNA can also be used to assess the effectiveness of treatment in real time (extensive review here (1)).

In the adjuvant setting, ctDNA has been used to predict recurrence. Patients who have detectable ctDNA, or minimal residual disease (MRD positive), after chemotherapy or surgery have an increased risk for recurrence compared to those who have no detectable ctDNA (MRD negative) (2,3). To understand the correlation of ctDNA dynamics and treatment effects on disease burden, longitudinal monitoring of ctDNA has also become a biomarker for patients undergoing treatment (4–12). In two studies of advanced stage patients with multiple solid tumor types being treated with immune checkpoint blockade, patients with a decrease or clearance of ctDNA while on treatment had improved progression-free survival (PFS) and overall survival (OS) compared to patients with an increase in ctDNA (5,11). Changes in ctDNA tend to precede imaging and may show treatment resistance, allowing for the adjustment of therapies (3,4).

To monitor ctDNA, next-generation sequencing (NGS) assays are categorized by variant composition: tumor-naïve or tumor-informed. Tumor-naïve monitoring utilizes fixed genomic targets, and panels capture a few variants from many patients (Supplemental Figure 1A). Tumor-naïve assays can be tailored to a specific tumor type (e.g. lung, colon, breast) or very broad (hundreds of genes) (13–16). Tumor-informed approaches rely on sequencing a pre-treatment sample and tracking a set of defined, patient-specific variants over time (9,17,18). The bespoke nature of the tumor-informed assays increases the assay design time, but sensitivity increases with the inclusion of more personalized variants compared to tumor-naïve assays. To overcome the smaller footprint of fixed gene sets and personalized panels, whole exome sequencing (WES) and whole genome sequencing (WGS) of liquid biopsy samples provides expanded breadth amenable to *de novo* variant discovery. Without the need for tissue or panel design, WES and WGS can be used for early detection of cancer, recurrence of disease, or treatment response (19,20). The expanded breadth of coverage usually results in lower sequencing depth, which impacts sensitivity for variant detection. Different bioinformatics strategies for tumor content analysis are being developed to boost sensitivity of the readout from WES or WGS (21).

This work presents a unique hybrid approach that enabled longitudinal monitoring of ctDNA in Gritstone’s GRANITE trial (NCT03639714), in which patients received an individualized neoantigen-directed immunotherapy. Each vaccine was comprised of twenty predicted neoantigens and was delivered using a heterologous prime-boost regimen comprised of chimpanzee adenoviral vector (ChAd) and self-amplifying mRNA (samRNA) vaccinations (22,23). The hybrid ctDNA assay targets two distinct sets of variants: an individualized set of variants from a pre-treatment biopsy and a common list of known oncogenic hotspots, recurrently mutated genes, and genes that play a role in the response to immunotherapy, including neoantigen-targeted therapies. The hybrid approach provided enhanced ability to monitor mechanisms of resistance in addition to the response to treatment.

## Results

### Tumor-informed and tumor-naïve ctDNA hybrid monitoring approach

For the twenty-nine patients treated, an average of 144 variants (range: 67-402) from the archival tissue biopsy were targeted for ctDNA monitoring (Figure 1A, Figure 1B, and Supplementary Table S1). By tissue type, the average was 114 variants (range: 67-192) for MSS-CRC patients (n=13), 149 (range: 76-241) for GEA patients (n=13), and 251 (range: 110-402) for NSCLC patients (n=3). Even though one MSS-CRC patient did not have whole blood collected for ctDNA monitoring, a personalized panel was designed for this patient (Supplementary Table S1). Hybrid capture baits were designed to capture all coding variants detected in WES of each patient’s archival tumor biopsy, which included the variants included in the patient’s individualized neoantigen-directed vaccine (neoantigen prediction described in (22,23)). Each set of patient probes was designed for a pool of multiple patients, resulting in a “pool of pools” of variants from six to nine patients (Figure 1C). Once a patient entered treatment with the vaccine regimen, longitudinal whole blood samples for ctDNA were collected approximately every four weeks with the patients’ scheduled dosing visits. An additional universal panel was designed, similar to tumor-naïve panels, to interrogate commonly mutated regions for *de novo* variant detection.

**Figure 1.**
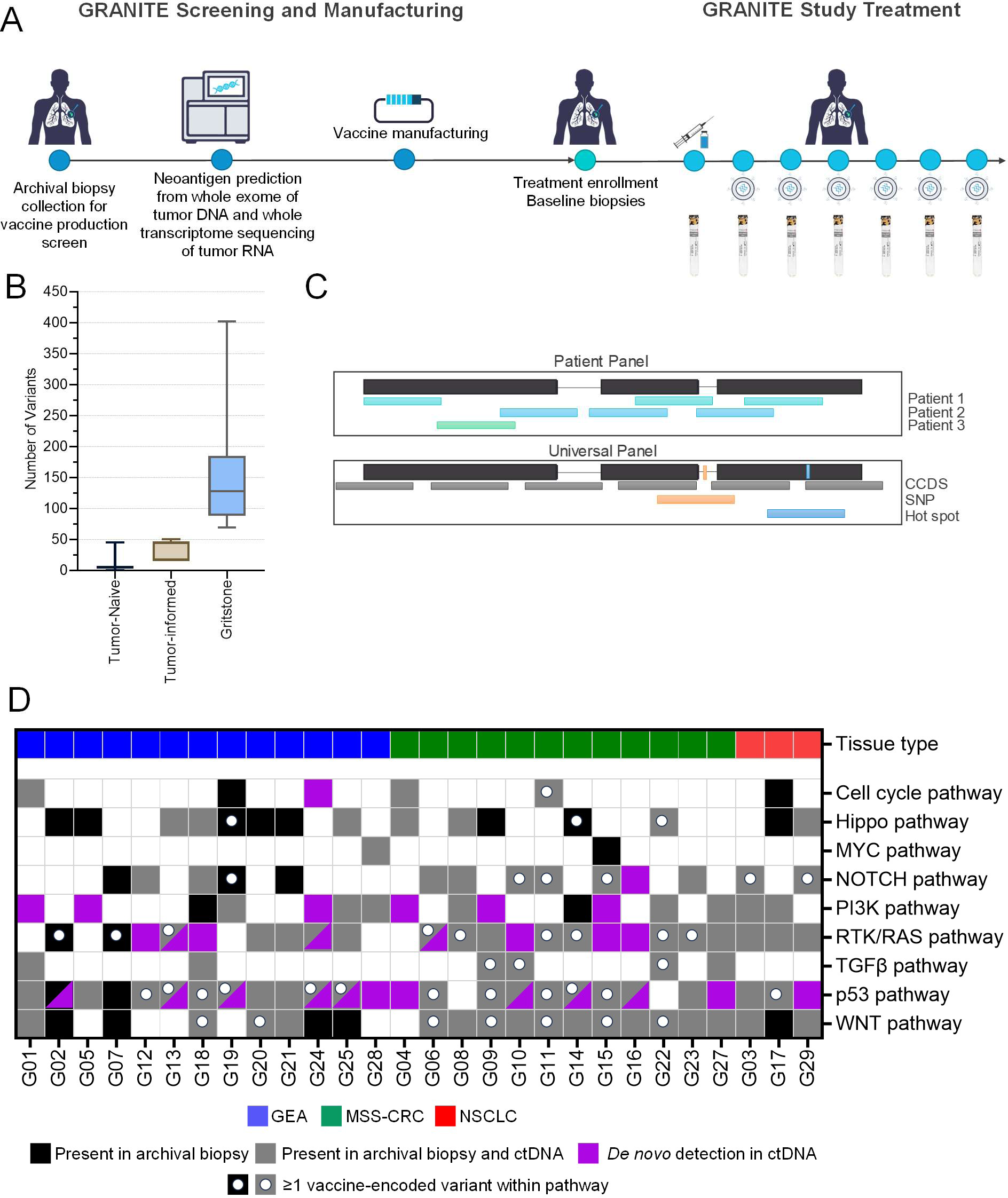
A) Whole exome sequencing of an archival biopsy was used for neoantigen prediction and the selection of variants for ctDNA monitoring. Upon enrollment in the treatment portion of the GRANITE study, blood draws for ctDNA were taken approximately every four weeks. B) In comparison to tumor naïve and other tumor-informed ctDNA assays, the ctDNA assay longitudinally monitors an average of 144 variants versus 6 (tumor-naïve) or up to 50 (tumor-informed). C) Each set of patient-specific targets is combined with other patients’ targets in a “pool of pools.” A universal panel was also designed to target SNPs and hot spots in all patients. D) The targeted ctDNA variants largely overlapped variants originally observed in the archival biopsies, including the vaccine-encoded variants, that were used for monitoring ctDNA dynamics.

The ctDNA assay leverages WES performed during the vaccine screening process, resulting in a substantial increase in the number of patient-specific tumor variants compared to the variants that would be monitored with curated tumor-naïve panels. Such panels cover an average of six variants per patient (Figure 1B and Supplementary Figure S1A) (13–15) and a median of one vaccine-encoded variant (range: 0-5 variants). The most prevalent vaccine-encoded variants were located within the p53 and RTK/RAS pathways, which are dominated specifically by *TP53* (11 unique variants in 12 patient neoantigen vaccines) and *KRAS* (3 unique variants in 6 patient neoantigen vaccines) (Figure 1D). Both genes are commonly interrogated in oncology panels but only account for 2.43% (14/576) of the unique vaccine-encoded neoantigens, demonstrating that the individualized neoantigens are mostly private to each patient. Using a tumor-naïve panel alone for monitoring patient-specific vaccine neoantigens limits the ability to observe any changes. Alternatively, options for commercial tumor-informed assays also utilize tumor biopsy WES, but only a maximum of fifty variants are targeted in those assays (24–26). Within those fifty, inclusion of the variants from the individualized neoantigen vaccine is not guaranteed; therefore, a comprehensive, individualized approach allows for not only monitoring patients’ tumor-specific neoantigens but also tracking of clonal and subclonal dynamics and heterogeneity (27,28). Furthermore, monitoring an expanded number of patient-specific variants improves the overall probability of detection, even at low variant allele frequency (VAF).

### Analytical characterization of the ctDNA monitoring assay

The monitoring assay was characterized using twenty-two mixtures of remnant patient cfDNA diluted into a background of cfDNA from healthy donors (Supplementary Table S2). Two patient-specific capture panels were designed and tested during characterization (see Methods). With the inclusion of patient SNPs diluted into a wild-type donor, variant calls were assessed for 9,911 variants at 2,301 positions (1,916 patient-specific and 385 germline variants) ranging from 0.02%-100% (Supplementary Figure S1B). The monitoring assay VAF values were concordant with whole exome sequencing (R^2^=0.983) with a limit of detection (LOD95) of 0.12% (Supplementary Figure S2A and Supplementary Figure S2B). For overall sensitivity and specificity, six healthy donors were sequenced using both patient-specific capture panels. Samples with an unknown variant status were also procured from five late-stage CRC and NSCLC patients and sequenced with a single patient panel. Overall, the assay sensitivity and specificity were 97.34% (CI: 97.03-97.66%) and 99.94% (CI: 99.93-99.95%), respectively, with an accuracy of 99.78% (CI: 99.76-99.81%) (Supplementary Table S3).

The ability to detect a change in the tumor content among ctDNA samples was also assessed. For each dilution, the entirety of the patient-specific variants, regardless of detection, were used to calculate the mean VAF, which ranged from 0.049-9.82% (R^2^=0.998), with most samples falling below 1% (Supplementary Figure S1C and Supplementary Figure S1D). Additionally, the variation of the measured mean VAF was assessed at different inputs and in replicates, including different operators, reagents, patient panels, and sequencing runs (Supplementary Figure S2C and Supplementary Figure S2D). The mean quantification for the lowest sample (Val005-01) measured 0.055% (Coefficient of variation (CV): 9.99%; n=4) and the next concentrated sample, Val005-02, which was double the concentration, measured 0.098% (CV: 8.33%, n=4), demonstrating the ability of the assay to detect small changes in ctDNA. Furthermore, the mean VAF was determined with different starting inputs of sample. For inputs of at least ten nanograms (ng), the measured mean VAF was consistent among replicates (CV: 0.3-5.7%), and for lower inputs of 2.5 and 5ng, the mean VAF values had more variability (CV: 6.0-20.8%). The values for all inputs ranged from 0.398-0.525% for patient panel 1 (orthogonal WES: 0.405%) and 0.172-0.289% for patient panel 2 (orthogonal WES: 0.221%).

The number of variants targeted by the assay buffers against large variability in the mean VAF, particularly with a lower input, where detection for a single variant is compromised by the limited number of molecules input into the assay (Supplementary Figure S3). By targeting numerous patient-specific variants, the overall sensitivity of the assay increases through increased likelihood of detecting any real variant. For individual variants approaching the assay LOD, using a single or very few variants confounds the ability to measure true changes in tumor burden and compare among different patient time points. Using the assay qualification samples, the mean VAF for each sample was calculated using random sets of *m* tumor variants and normalized to the expected mean VAF from orthogonal WES (Supplementary Figure S3A). With only a single random variant, the calculated mean VAF could range from 0x (not detected) to 2x of the expected value. With 50 tumor variants, the distribution narrows to 0.8-1.2x of the mean VAF. To further understand the variability of the mean VAF, the CV of the mean VAF was calculated among sets of technical replicates. Using a single variant, the CV ranged from 1-10%, and any combination of 50 variants resulted in a mean VAF with a CV of less than 2%. The larger number of variants protects against any one region or variant substantially affecting the mean VAF due to factors that may compromise detection, such as sequencing depth (pseudogenes, off-target coverage, input molecules) or low VAF (near or below LOD) (Supplementary Figure S3B).

### Monitoring of tumor-informed variants

Patient-specific panels in the monitoring assay were designed to capture the vaccine-encoded variants, which would not usually be captured even by bespoke commercial panels, and other tumor tissue variants to understand tumor evolution under the vaccine regimen. To achieve the sensitivity often required for ctDNA detection, the longitudinal monitoring assay leverages duplex sequencing for error correction (29). cfDNA libraries were sequenced to an average raw mean paired-end depth of 169,776x, which was reduced to 4,868x after duplex collapse (Supplementary Table S4). Paired gDNA from whole blood or buffy coat was sequenced to 137,355x for an average duplex depth of 3,815x. For patients with biopsies of high-quality DNA without formalin-fixation, the DNA remains intact, and a high depth of duplex sequencing can be achieved for variant calling to as low as approximately 1%. For patients with available biopsies, the libraries were sequenced to a mean paired end read depth of 129,248x for a duplex depth of 3,089x.

Because patients received multiple lines of therapy prior to treatment with the individualized neoantigen-directed immunotherapy, variants may have changed through prior courses of treatment and tumor evolution. cfDNA was collected for 12/13 MSS-CRC patients (92%), and the number of vaccine-encoded variants detected across the longitudinal samples (number of samples: 1-17, average: 8) ranged from 13-20 (65-100%; average: 85%) (Figure 2A). For GEA patients, 2/13 (15%), G02 and G07, had no evidence of disease post-resection, and no targeted variants were detected in their cfDNA. Of the remaining 11 GEA patients with 1-18 longitudinal cfDNA samples (average number of samples: 6), 11-20 (55-100%, average: 16) of the vaccine-encoded variants were detected. In the NSCLC cohort, 3, 5, and 8 cfDNA samples were collected. All twenty vaccine-encoded variants were detected in 2/3 patients, and 9 vaccine variants (45%) were detected in the third patient. For the vaccine-encoded variants that were not detected in ctDNA, many of them appeared to be subclonal variants in the archival biopsy (Supplementary Figure S4A).

**Figure 2.**
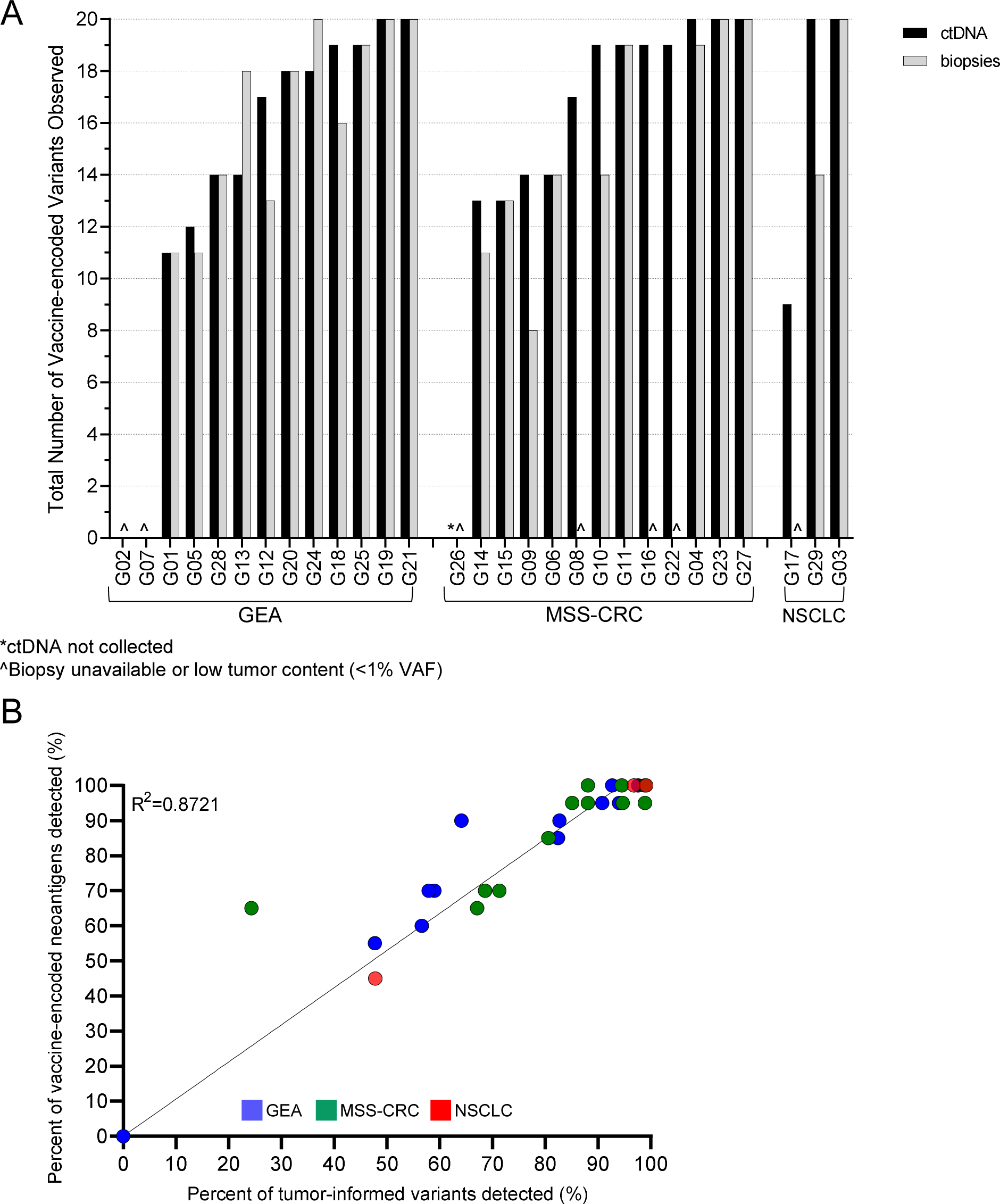
Variant detection from duplex sequencing of cfDNA and biopsies. A) The majority of vaccine-encoded variants were detected in both ctDNA and biopsies in most patients where sample was available. B) The percentage of vaccine-ended neoantigens largely correlated with the detection of all the variants that were targeted by the ctDNA assay.

Leveraging the sensitivity of duplex sequencing, a total of 29 patient biopsies from 23/29 (79%) patients across the three tissue types were sequenced with the monitoring assay (Supplementary Table S1). Most patients had only a single biopsy available (17/23; 74%), and six had two tissue biopsies available for duplex sequencing. One of the patient biopsies (1/29; 3.4%) could not be assessed due to limited tumor content (<1% VAF). In the MSS-CRC patient biopsies, 8-20 (40-100%, average: 75%) vaccine-encoded variants were found in tissue, and in the GEA patients, 11-20 (55-100%, average: 80%) vaccine-encoded variants were detected. In the two NSCLC patients with available biopsies, 14/20 (70%) and 20/20 of the vaccine-encoded variants were detected (Figure 2A).

Considering all captured tumor variants, the percentage of vaccine-encoded variants largely coordinated with the detected non-vaccine tumor variants in ctDNA (Figure 2B and Supplementary Figure S4B). In the MSS-CRC cohort, three cfDNA samples and a single tissue biopsy were collected for patient G15, who was the patient with the highest overall variant loss when compared with the archival biopsy. Of the 259 targeted variants (including the 20 vaccine-encoded variants), 13/20 (65%) of the vaccine-encoded variants and 50/239 (21%) of the non-vaccine variants were observed in both the ctDNA (average VAF: 37-40.5%) and the tissue biopsy (average VAF: 1.90%) (Supplementary Figure S5A and Supplementary Figure S5B). Both the liquid and tissue biopsies demonstrated loss of the same variants (Supplementary Figure S3C). For the remaining 25 patients with ctDNA-positive samples, 47-100% (average: 79.2%) of the non-vaccine tumor variants and an average of 17/20 (85%) vaccine-encoded variants were detected in ctDNA. In the corresponding tissue samples, 42-100% (average 75%) of non-vaccine variants and 40-100% (average: 80%) of vaccine-encoded variants were detected (Supplementary Figure S4B). Even with neoantigen and variant loss, the original neoantigen prediction from the archival tumors resulted in neoantigens that could still be targeted using the individualized neoantigen vaccine, and the patient-specific nature of the assay allowed for detection in both liquid and tissue biopsies.

Where tissue was insufficient, collecting longitudinal cfDNA samples provided tumor information with a regular cadence and was often more comprehensive than a single tissue biopsy. The ctDNA provided a better reservoir than the tissue biopsies of the tumor variants in all but two patients, G13 and G24, both of whom were GEA patients (Figure 2A, Figure 3A, and Supplementary Figure S5D). G13 had no detectable ctDNA variants until 20 weeks after starting treatment, which peaked at 0.6% VAF at week 40 (Supplementary Figure S5C). A baseline tissue biopsy was not collected for this patient. An on-treatment tissue biopsy contained 18/20 (90%) of the vaccine-encoded variants and 49/56 (87.5%) of the archival tumor tissue variants whereas 14/20 (70%) of the vaccine variants and 30/50 (60%) of the archival tumor tissue variants were detected in ctDNA (Supplementary Figure S5D). The overall cfDNA level of this patient was low (median cfDNA yield: 8.39 ng/mL plasma vs 14.9 ng/mL for GEA patient samples and 15.8ng/mL for all samples); therefore, the lack of detection could be attributed to low tumor shedding or very low tumor content.

**Figure 3.**
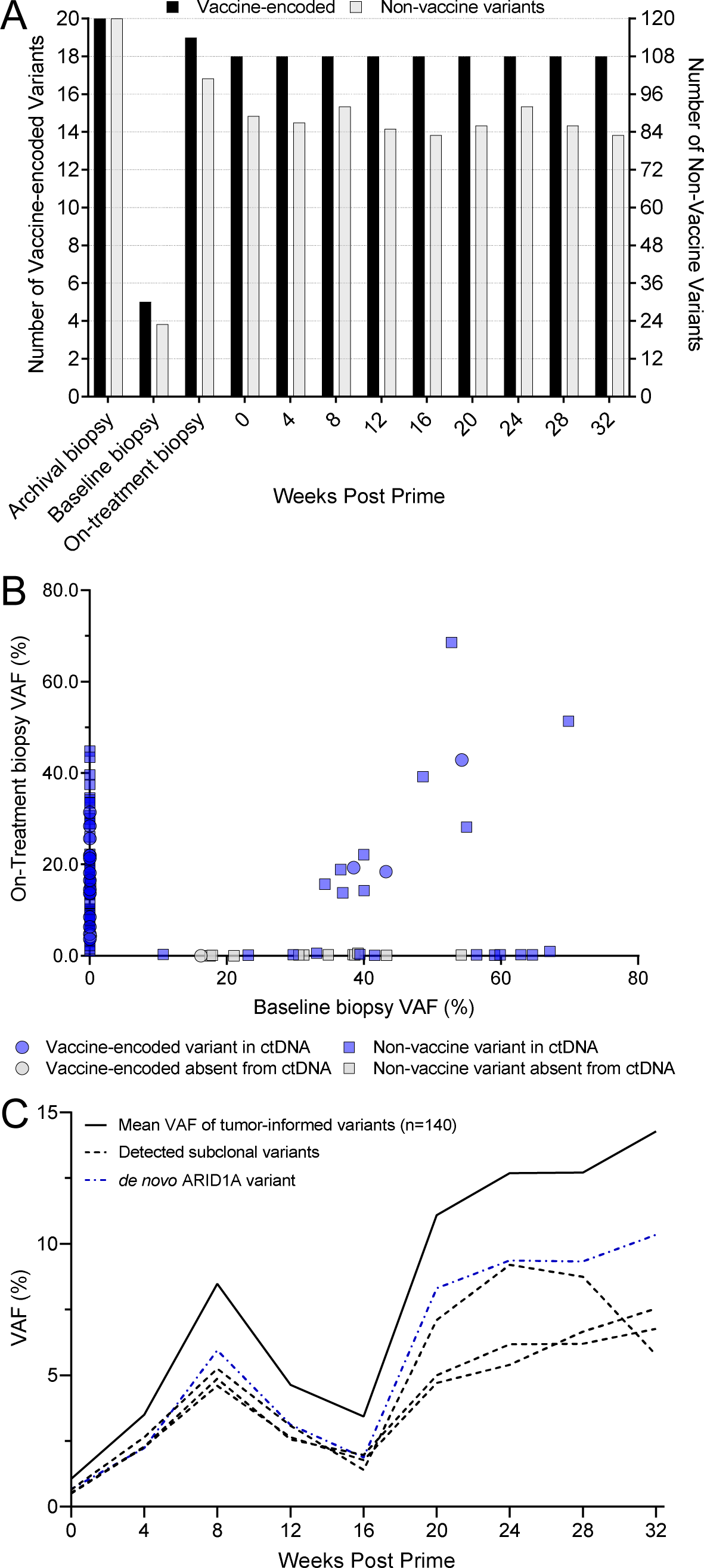
G24 variant detection in ctDNA and biopsies A) Longitudinal variant detection in biopsies and cfDNA of vaccine-encoded and non-vaccine variants indicate similar detection levels among timepoints. B) Comparison of variant detection in ctDNA and biopsies. The lack of variant overlap among multiple biopsies and ctDNA demonstrates tumor heterogeneity within the primary tumor site that was not fully captured by ctDNA. Even in tumor-informed assays, heterogeneity poses a challenge to accurately monitor response. C) Longitudinal ctDNA in patient G24. The dynamics of 140 targeted variants was closely followed by subclonal tumor variants and a *de novo* variant found in *ARID1A*.

For the GEA patient G24, monitoring for 139 variants from the archival tumor tissue was performed on nine cfDNA samples and two tissue biopsies. Of the 139 targeted tumor variants, 19 (13.7%) were not found in any of the ctDNA samples or tissue biopsies (Figure 3A). Despite both biopsies being collected from the primary tumor site, the targeted tissue biopsy variants had little overlap, and the on-treatment tissue biopsy more closely resembled the archival tissue biopsy in variant composition (Figure 3A and Figure 3B). Among the targeted variants, only twelve variants, three being vaccine-encoded, were shared between the baseline and on-treatment tissue biopsies (Figure 3B). The other targeted variants found in the baseline tissue biopsy were either undetected in ctDNA or detected at frequencies below 0.25% in only one or two time points.

For each patient design, an additional set of tumor-informed variants from the archival biopsy was included in the patient-specific panel for exploratory clonal evolution. The additional tumor-informed variants were chosen from those below the VAF threshold for neoantigen prediction but had high mapping and base qualities. Ten of the exploratory variants in patient G24 were present at high VAF (10.7-54.2%) in the baseline tissue biopsy and were mostly absent from the patient’s ctDNA and on-treatment tissue biopsy. Another four of the exploratory variants were present only in the on-treatment tissue biopsy and three of those four followed the overall cfDNA trend (Figure 3C and Supplementary Table S5). Patient G24 had a heterogeneous tumor, and the baseline tissue sample had variants that were primarily at the lowest VAF in the archival biopsy (Supplementary Table S4). With only a few of the exploratory variants from the baseline tissue biopsy having been present in ctDNA at low frequency, the lack of representation in ctDNA could indicate different portions or rates of the tumor shedding, extensive heterogeneity (diversity of variants not captured), or a changing tumor landscape.

To further elucidate the heterogeneity observed in the liquid and tissue biopsies of G24, beta-allele frequency (BAF) was analyzed at the patient’s heterozygous SNP positions covered by the ctDNA assay. The minor allele positions show changes in the cfDNA over the course of treatment and differences between the two tissue biopsies for G24 (Supplementary Figure S6). From the duplex sequencing, which collapses reads to single molecules, the coverage at the SNP rs879732 ranges from 59,654x in the baseline cfDNA (mean coverage: 7,410x) to 484,282x (mean coverage: 6,232x) in the final sample (32 weeks post prime). The reference allele drops to less than 0.79% in the cfDNA at 32 weeks post prime whereas the matched blood gDNA indicated heterozygosity (46% VAF with a duplex coverage of 2,266x). The loss of heterozygosity (LOH) and amplification were not observed in G24’s baseline tissue biopsy: 47% VAF with 4,579x coverage (mean coverage: 3,532x). Even though it was not captured in the baseline biopsy, because the LOH was initially captured in the baseline cfDNA sample and ultimately in the on-treatment tissue biopsy (98% VAF), a portion of the tumor harbored the amplification prior to treatment. Furthermore, the longitudinal BAF followed the tumor variant ctDNA dynamics: an expansion occurred at 8 weeks post prime, followed by contraction at week 16, and further expansion through the end of treatment at 32 weeks. With over 100 variants used for monitoring, including subclonal variants, and the additional SNPs across the genome, the breadth of the assay supported a broader understanding of the tumor composition, and it also highlighted the discrepancies that persist between liquid and tissue biopsies over the course of a patient’s disease.

### Tumor-Naïve Variant Detection

In addition to the patient-specific tumor variants, a set of genomic regions were included in a universal panel for monitoring in all patient samples. The panel covered commonly mutated genes (e.g., *TP53*, *ARID1A*), hot spots (*KRAS* G12 and G13 codons) and genes involved in antigen presentation or resistance in immunotherapy (*B2M*, *TAP1*, *TAP2*) (30–34). Tracking tumor-naïve variants may provide evidence of resistance to treatment, heterogeneity, or additional *de novo* variants that were either acquired or originated from additional tumor sites not represented in the patient-specific variants. Biological noise from clonal hematopoiesis (CH) can contribute to false positive somatic calls in the sequencing of cfDNA (35–37). Paired sequencing of DNA from buffy coat, whole blood, or PBMCs was used to remove suspected CH variants (Supplementary Figure S7D). Among the 28 patients with cfDNA samples, 21 (75%) had a *de novo* variant detected in a region captured by the universal panel (Figure 4A).

**Figure 4.**
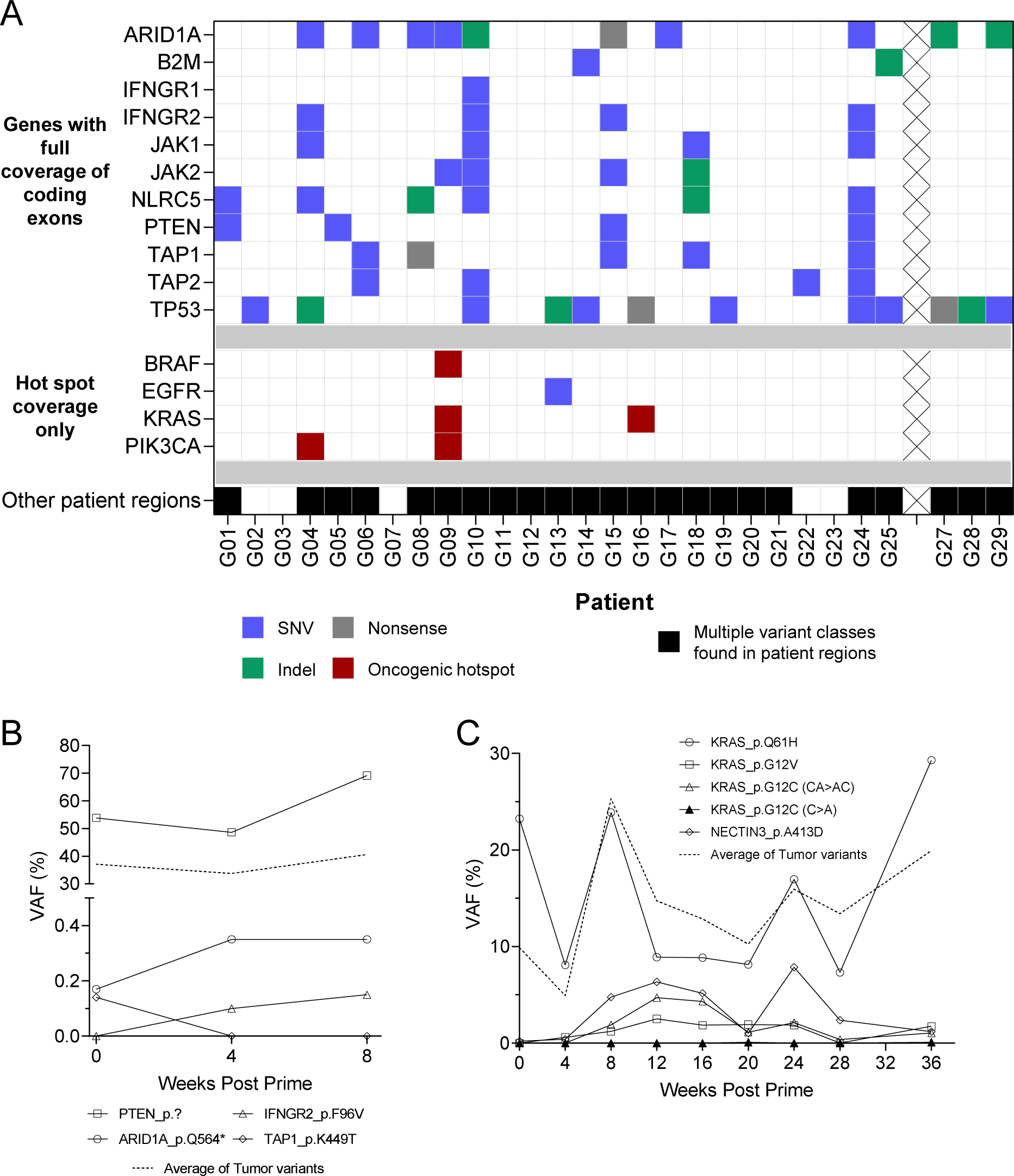
*De novo* variant detection in ctDNA. A) *De novo* variants detected in ctDNA in tumor-naïve regions included in the monitoring assay. B) G15 ctDNA dynamics of *de novo* variants. Although most *de novo* variants were at a low frequency, some were found to correlate closely with tumor-informed ctDNA dynamics. C) G09 ctDNA dynamics of *de novo* variants that included multiple KRAS variants.

In 25 patients with *de novo* variants, many of them were detected at a low VAF in a single cfDNA timepoint (Supplementary Table S6), but in select patients, multiple variants were consistently detected in longitudinal cfDNA samples (Figure 4B, Figure 4C, and Supplementary Table S5). Patient G15 acquired a splice variant in *PTEN* at a similar VAF to the patient-specific tumor variants, and variants in *ARID1A*, *IFNGR2*, and *TAP1* appeared at allele frequencies below 0.5% (Figure 4B and Supplementary Table S6). In patient G09, multiple *KRAS* variants were detected that had not been detected in either the archival biopsy or the follow-up biopsy. A *KRAS* Q61H variant in this patient followed the same trajectory as the archival tumor variants whereas three new *KRAS* G12 variants occurred at a lower VAF with different dynamics (Figure 4C and Supplementary Figure S7A). Although the variants tended to change similarly in VAF over time, separate populations of variants were observed in G09 (Supplementary Figure S7B). The *KRAS* Q61H variant in G09, which tracked with the mean VAF dynamics, was likely present in the tumor but not represented in the biopsied tissue (Figure 4C and Supplementary Figure S7B). Alternatively, the variant may have been acquired in the time between the archival biopsy collection and the start of treatment. The multiple KRAS G12 hot spot variants also present in this patient demonstrate the genomic changes that can occur in advanced disease and the utility of ctDNA to capture those changes.

In patient G08, variants in antigen presentation genes were detected in the ctDNA, but because the tumor tissue biopsy content was too low, they were not detected in the tissue. Notably, at the beginning of treatment, a frameshift variant was detected in *NLRC5*, the transactivator of HLA class I, and the frameshift variant followed the mean VAF of the targeted patient-specific variants over the course of treatment. This patient showed a substantial drop in ctDNA at weeks 28 and 32 before a rebound in ctDNA at 36 weeks (Supplementary Figure S7C and Supplementary Table S5). At 40 weeks past the vaccination priming dose, a new missense variant in *NLRC5* was detected, and at 57 and 61 weeks, two *TAP1* variants, one a frameshift and another a nonsense variant, emerged. All four variants (two in *NLRC5* and two in *TAP1*) were consistently detected in the ctDNA through the end of treatment at 65 weeks. *TAP1*, with *TAP2*, is involved in the loading of peptides into the MHC complex for presentation on the cell surface (32,38–40). The loss of function variants in *TAP1* would downregulate the presentation of tumor-specific neoantigens, which would suggest possible resistance to the immune pressure elicited by the vaccine. Analysis of this patient’s ctDNA using the ability to detect both patient-specific variants and emerging variants demonstrates the value of ctDNA to be a valuable and accessible biomarker to assess treatment response and disease advancement.

### Loss of heterozygosity from ctDNA

The universal panel included the capture of the coding regions of the MHC class I genes: *HLA-A*, *HLA-B*, and *HLA-C*. The sequencing reads from these genes were extracted after initial alignment and mapped onto a patient-specific reference of their HLA alleles: a method that has been used previously for tissue sequencing (41,42). The sequencing depth and inclusion of duplex unique molecular identifiers (UMIs) provided the resolution to assess LOH in not only the tissue biopsies but also longitudinal cfDNA samples. Analysis of LOH at the HLA loci gives insight into an additional avenue of immune escape and tumor-intrinsic resistance to immunotherapy; surface presentation of some neoantigens would be reduced with the loss of the presenting HLA allele.

One GEA patient (1/13; 7.7%), G19, and two MSS-CRC patients (2/12; 16%), G04 and G23, demonstrated full haplotype loss, which had been present in the archival biopsies (Figure 5A and Figure 5B). Using the monitoring assay, the haplotype loss was observed in the follow-up tissue biopsies for all three patients. LOH was not observed in the ctDNA sample from G19 despite being observed in the archival and baseline tissue biopsies. In MSS-CRC patient G04, who was homozygous in *HLA-C*, a private SNP within HLA-C*07:01 demonstrated a change in allele ratios. Using G04’s private, heterozygous SNPs near the HLA loci along chr6 also covered by the assay, including within *TAP2*, the ratios of the different alleles showed a similar imbalance (Supplementary Figure S8A). Combining the LOH observed in *HLA-A* and *HLA-B* in the same haplotype and LOH in SNPs both downstream and upstream of the HLA region on chr6, the genomic change was likely a larger event affecting class I, class II, and the TAP genes. Similarly, G23 had full haplotype loss within the class I genes and LOH within *TAP2* (Supplementary Figure S8B). Chromosomal instability was not isolated to the HLA locus: both G04 and G23 had evidence of aneuploidy in the baseline tissue biopsies and ctDNA samples (Figure 5C and Figure 5D). Loss of heterozygosity across the MHC and antigen presentation genes may allow the tumor to evade immunity through reduced or altered presentation of immunogenic antigens. Changes in TAP expression have been found to downregulate MHC class I expression and present new peptides in a TAP-independent manner (43,44).

**Figure 5.**
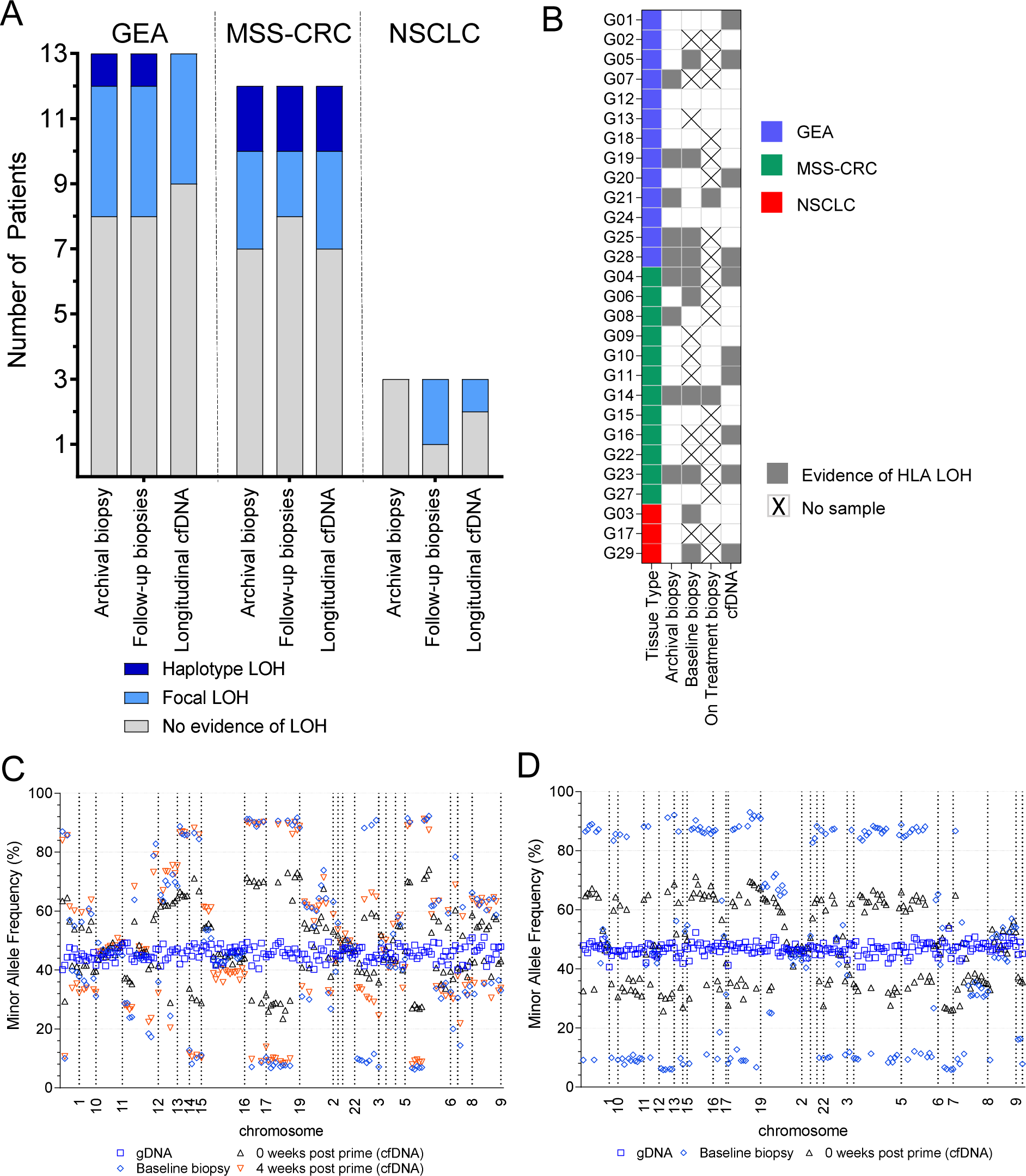
Evidence of HLA loss of heterozygosity (LOH). A) HLA LOH by tissue type. Biopsies and cfDNA were analyzed for evidence of HLA LOH. B) HLA LOH by patient. HLA LOH evidence by patient and sample. C) Using B-allele frequency of 184 heterozygous SNPs, patient G04 showed genomic instability beyond HLA LOH that was present in both cfDNA and the baseline tissue biopsy. D) In patient G23, B-allele frequency analysis of 204 heterozygous SNPs indicated genomic instability in the baseline biopsy and cfDNA. Patients G04 and G23 were MSS-CRC patients with evidence of full haplotype loss.

Aside from full haplotype LOH, focal LOH in either a mixture of the HLA genes (e.g. *HLA-A* and/or *HLA-C* only) or across the alleles (e.g., *HLA-A* and *HLA-C* from one haplotype and *HLA-B* from the other haplotype) was also observed. An additional 7/13 (54%) GEA patients had focal LOH in one or more loci (Figure 5). In the MSS-CRC cohort, 6/12 (50%) patients had focal HLA LOH. Among the NSCLC patients, the baseline tissue biopsies collected from two of the patients (2/3; 67%) showed evidence of focal HLA LOH, but only patient G29’s final ctDNA sample showed HLA LOH (Figure 5B).

Although most patients with HLA LOH had evidence of it prior to the initiation of the neoantigen immunotherapy, two patients demonstrated LOH while being on treatment. The first treated patient (G01), a GEA patient who was on study for approximately 18 months, had 237 targeted variants from the archival biopsy, and 113/237 (48%) were detected in ctDNA. Using 102 heterozygous SNP positions included in the assay, this patient’s cfDNA indicated SNP divergence starting at week 24, corresponding to a sharp increase in ctDNA (Figure 6A and Figure 6B). In the final ctDNA sample, some of the tumor variants reached 80-95% VAF, which could be partially attributed to aneuploidy within the tumor. Evidence of LOH was found for *HLA-A*, *B*, and *C* at 44 weeks, 72 weeks, and in the final sample at 80 weeks post prime, which correlated with the ctDNA tumor dynamics and BAF analysis of the patient’s heterozygous SNPs (Figure 6C). Evidence of LOH in *HLA-C* was also observed at 40 weeks and again at 48 weeks. The patient received a second dose of individualized chimpanzee adenovirus (ChAd) at week 72 that was followed by a sharp decrease in ctDNA until the final increase in ctDNA at 80 weeks. The LOH observations were focal imbalances rather than a full haplotype loss; *HLA-A* and *HLA-C* from one haplotype were lost, and *HLA-B* from the second haplotype was lost. The onset of LOH may indicate immune pressure exerted by the neoantigen vaccine and, with the presence of HLA LOH, the change in antigen presentation would be expected to assist in immune escape (32,45).

**Figure 6.**
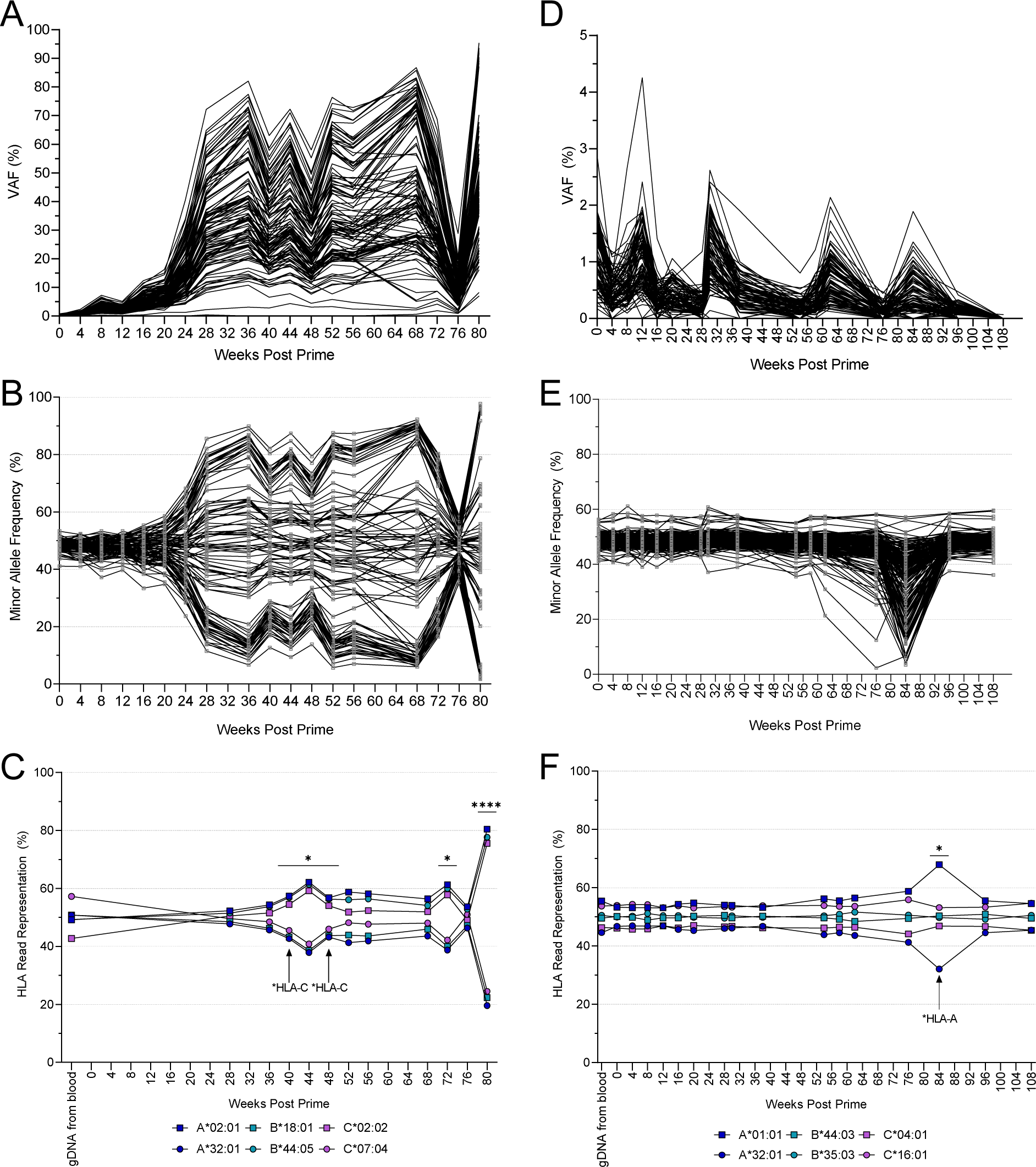
GEA patient G01 and MSS-CRC patient G16 longitudinal ctDNA dynamics. A) ctDNA dynamics of 113 (out of 237) patient-specific variants of patient G01 B) B-allele frequency of 102 SNPs tracked in the cfDNA of patient G01 showed aneuploidy and LOH. C) HLA allele representation in the cfDNA of patient G01. HLA LOH was initially observed at 40 weeks after initial vaccination. D) Dynamics of 90 (out of 96) targeted variants detected in the ctDNA of G16. In the final timepoint, only a single variant was detected. E) Longitudinal monitoring of 190 SNPs in cfDNA indicated LOH across the genome at 84 weeks. F) HLA allele representation in cfDNA of patient G16 showed evidence of HLA LOH only at week 84, correlating with the LOH observed at SNP positions genome-wide. *p<0.05, ****p<0.0001

To further elucidate the instability observed in G01’s ctDNA, residual cfDNA material (shotgun libraries or nucleic acid) from G01 underwent whole exome sequencing (WES) with duplex UMIs, which were used to create consensus of any PCR duplicates. Because the error correction is less stringent and the coverage depth is reduced, variant detection is reduced, but the WES dynamics mirrored that of the monitoring assay without the granularity afforded by sequencing all available cfDNA samples (Supplementary Figure S9A). HLA LOH was only evident at 80 weeks, which may be attributed to achieving half the sequencing depth from the increased footprint of WES. Additional heterozygous positions within the exome were captured, showing the spread of allele frequencies on chromosome 6 to support the HLA imbalance and a sharp separation of alleles on chr9 (Supplementary Figure S9C). Located on chromosome 9 is *JAK2*, a downstream target of interferon γ signaling as part of the JAK-STAT pathway, and changes in sensitivity to *IFN-γ* may confer treatment resistance even with T-cell infiltration in the tumor microenvironment (45–47).

MSS-CRC patient G16 showed evidence of HLA LOH in ctDNA (Figure 6D, Figure 6E, and Figure 6F). At 84 weeks post initial vaccination, evidence of LOH was observed in the ctDNA in *HLA-A,* but it was not seen in any of the other ctDNA samples from the patient either before or after the sample at 84 weeks. The patient had liver and lung lesions and was found to have brain metastasis. Even with relatively low tumor content (<1%), a distinct change in the BAF was seen in the ctDNA, which supports the HLA LOH observation (Figure 6E and Figure 6F). A possible explanation for the sudden changes at week 84 could be metastatic sites with little tumor shedding previously or a lack of variant representation in the ctDNA panel. This evolution was instead captured in the ctDNA using a combination of the patient’s private SNPs and interrogation of the HLA class I loci. The patient nearly achieved clearance of ctDNA by the end of treatment; in the final sample at 108 weeks, only a single tumor variant was detected in the ctDNA at 0.07% (Figure 6D).

Importantly, both patient G01 and G16 lacked high quality tissue biopsies for sequencing in parallel with the ctDNA. Patient G01’s initial on-treatment tissue biopsy consisted of little to no tumor content, and the two later tissue biopsies were formalin-fixed and paraffin embedded (FFPE). The final biopsy collection near the end of G01’s treatment originated from a brain metastasis that underwent WES, and it was the only tissue sample from this patient with evidence of HLA LOH (Supplementary Figure S10A). Patient G16’s brain metastasis was resected and underwent WES, but the tissue sample from the resection did not have any evidence of HLA LOH (Supplementary Figure S10B). These patients further exhibited the limitations of tissue biopsies that can be overcome with longitudinal ctDNA monitoring.

## Discussion

The insights gained from liquid biopsies have been recognized for their utility, specifically for precision medicine (3,48). Despite the utility of ctDNA, studies will be required to understand the relationship between ctDNA dynamics and changes in solid tumor tissue encountering various treatment modalities (chemotherapy, TKI, CPI, or combination thereof) (5,9,10,49,50). For patients receiving GRANITE, an individualized neoantigen-directed immunotherapy, ctDNA monitoring necessitates a comprehensive, individualized approach to ensure successful monitoring of the vaccine-encoded variants and the tumor evolution during treatment.

The monitoring assay demonstrates high sensitivity and specificity (>95%) with low technical variability to allow for accurate quantification of ctDNA over time. Because the assay targets most of the variants found from WES of the tumor tissue, the assay also has increased sensitivity for samples with a low tumor content or low ctDNA shedding that would be compromised in assays targeting fewer variants. Although the assay was designed specifically for cfDNA, matched gDNA was used to remove variants from clonal hematopoiesis, background artifacts, and SNP identification. Highly intact DNA from biopsies were also sequenced using duplex sequencing, which identified variants that would be missed in many of the study biopsies with low tumor content (<10%) when analyzed in workflows without UMIs. The liquid and tissue biopsies showed strong concordance of the targeted patient-specific variants (>95%), with ctDNA harboring more of the variants than tissue biopsies in all but two patients. During the design of the ctDNA monitoring assay, the tissue source and heterogeneity of the tumor biopsy could affect the variants selected for monitoring. Tumor heterogeneity not immediately evident in a tissue biopsy could enrich for variants originating from a subclonal portion of the tumor. The inclusion of subclonal variants for monitoring attempted to mitigate the bias heterogeneity can induce, which was revealed, for example, in patient G24. The individualized design from an archival tissue biopsy alone, however, may not detect new variants that could be detected with static gene panels, whole exome, or whole genome sequencing, preventing a more comprehensive view of clonal evolution where primary and metastatic sites diverge.

The heterologous prime/boost vaccine regimen in GRANITE study NCT03639714, a chimpanzee adenovirus (ChAd) prime vaccination followed by self-amplifying mRNA (samRNA) boost vaccination, showed a unique ctDNA dynamic in select patients. In the samples 4-8 weeks post-prime, a transient increase of ctDNA (“molecular pseudoprogression”) was observed that would then recede by 12-16 weeks after the prime vaccination, which highlights the need for an individualized assay to understand the dynamics and mechanism of a unique vaccine platform. The increase was observed in all three tumor types and would thus be specific to treatment with the neoantigen-directed immunotherapy post the ChAd prime. The ctDNA dynamics and treatment response will be explored further in a randomized Phase 2/3 trial where GRANITE is administered in the first line setting for MSS-CRC patients (NCT05141721).

As demonstrated by the detection of *de novo* variants in 75% of the patients, including drivers (*KRAS* and *BRAF*) and resistance variants (*TAP1*), the hybrid approach of using tumor-informed and tumor-naïve panels expands the utility of the ctDNA monitoring assay. The universal panel was designed to include common hot spots, including full coding coverage of select genes. Multiple universal panels could be designed to specific tissue types for more versatility. In addition to hot spots, the universal panel targeted common SNPs that were used for both LOH analysis and the detection of aneuploidy.

For a neoantigen-directed vaccine, antigen presentation is critical for engagement with T-cells, and the identification of tumor-intrinsic perturbations provides an avenue to correlate and understand response to treatment. Accordingly, the ctDNA monitoring assay provided insights into the tumor changes and resistance mechanisms observed in many of the patients, including acquired aneuploidy, HLA LOH, and additional variants in oncogenes or tumor suppressors. Of the cohort, 36% of patients showed evidence of LOH in at least one of the canonical HLA class I genes in ctDNA. All except two of the ten patients with LOH presented HLA LOH prior to the initiation of vaccine treatment. Because HLA LOH may affect the diversity of neoantigens presented for T-cell interaction, monitoring regions outside the tumor-specific variants elucidates potential changes in response to the treatment.

The individualized design and expanded longitudinal ctDNA monitoring demonstrated the granularity and type of genomic information that can be gained from such analyses of longitudinal ctDNA and will continue to be critical to the field of precision medicine. The ability to detect ctDNA with high sensitivity (< 0.1% tumor content) and high specificity allows for a more robust assessment of ctDNA content, which is crucial for robustly quantifying minimal residual disease (MRD) and ctDNA dynamics. With a more amenable and easier liquid biopsy collection, the breadth of the assay enables effective use of ctDNA monitoring for discovery and understanding of treatment effect on tumor evolution that includes control and/or progression of disease and potentially emerging resistance. Such an assay, therefore, will be critical to the development of innovative cancer therapies with a potential to benefit patients across tumor types.

## Acknowledgements

The authors would like to thank the patients and their families. Funding was provided by Gritstone bio. Employees of Gritstone bio received salaries for their work.

## Materials and Methods

### Patient Consent and Enrollment

The clinical study and all related analyses were carried out in accordance with the Declaration of Helsinki and Good Clinical Practice guidelines and was approved by the appropriate institutional review board (IRB) or ethics committees at each participating site. All patients provided written, informed consent.

Patients gave initial consent for the screening portion of the GRANITE GO-004 (https://clinicaltrials.gov/study/NCT03639714) study for eligibility screening, as described previously (23). Briefly, an archival biopsy in either FFPE block or slides was sent to Gritstone bio for assessment of tumor content via H&E prior to sequencing. Whole blood was collected in a K_2_EDTA tube and sent to Gritstone for extraction of matched normal gDNA from whole blood. Neoantigen prediction was performed from the sequencing of the tumor DNA and RNA. Prior to treatment with the individualized chimpanzee adenovirus and self-amplifying mRNA, patients provided a separate consent for the study treatment portion of the study.

### Archival Biopsy Processing

Tissue with at least 10% tumor content via pathology review had DNA and RNA extracted using the Qiagen AllPrep FFPE DNA/RNA kit (Qiagen, Waltham, MA). gDNA from 200uL whole blood was extracted using the Qiagen QIAamp Mini Blood kit. Prior to library preparation, gDNA and tumor DNA were sheared to an average fragment length of 200bp using a Covaris ultrasonicator (Covaris, Woburn, MA).

Libraries for whole exome sequencing (WES) of matched normal gDNA and tumor DNA and whole transcriptome sequencing (WTS) of tumor RNA were prepared using NEBNext UltraII reagents (New England Biolabs, Ipswich, MA). Shotgun libraries were captured overnight using the IDT exome panel with IDT hybridization reagents (Integrated DNA Technologies, Coralville, IA). Captured libraries from a single patient (gDNA, tumor DNA, and RNA) were pooled at a 1:4:1 ratio, respectively, and sequenced across three lanes of a HiSeq or pooled for sequencing on a NovaSeq S1 using 2×75 (HiSeq) or 2×101 (NovaSeq) read lengths (Illumina, San Diego, CA). Variant calling and neoantigen prediction were performed as described previously (22,23).

### Follow-up Biopsy Collection and Extraction

For patients who initially passed the EDGE neoantigen burden (ENB) threshold and were eligible for study treatment, patient consent was obtained for study treatment. When amenable, a baseline biopsy was collected approximately four weeks prior to the start of treatment. Core needle biopsies were fixed in RNALater (ThermoFisher, Waltham, MA) or flash frozen in liquid nitrogen at clinical sites before being shipped to Gritstone bio. DNA and RNA from biopsies was extracted from the tissue using the Qiagen AllPrep DNA/RNA kit with a proteinase K digestion step. gDNA was quantified using a Qubit fluorometer, and quality was checked on TapeStation Genomic DNA (Agilent, Santa Clara, CA).

### Matched Normal gDNA

Matched normal genomic DNA for each patient was obtained from either the extraction of whole blood that had been collected in K_2_EDTA tubes or the extraction of PBMCs. PBMCs were isolated using a density gradient and 50,000 cells were isolated for the extraction of gDNA using the Qiagen AllPrep DNA/RNA kit.

### cfDNA collection

Whole blood was collected in Streck cfDNA blood collection tubes (BCT) (Streck, Omaha, NE). For patient samples, two tubes of whole blood were collected and processed immediately at sites using a double-spin protocol. Whole blood was centrifuged at 1600xg for 10 minutes. The plasma layer was removed and spun at 5000xg for 10 minutes to remove any residual cellular debris. The plasma was removed from any debris and stored at -80°C. For samples that could not be processed at clinical sites due to COVID restrictions, the whole blood was shipped overnight to Gritstone and processed with a double-spin protocol upon receipt.

Whole blood from healthy donors was collected in Streck cfDNA BCT. Upon collection, the whole blood was either delivered the same day for processing at Gritstone or shipped overnight for processing the next day. Whole blood was centrifuged at 1600xg for 10minutes. The plasma layer was removed and spun at 5000xg for 10 minutes. After the second spin, the plasma was removed from any debris and stored at -80°C until extraction. The buffy coat layer was removed and stored at -80°C to be used for the extraction of matched normal gDNA. gDNA was extracted using the Qiagen DNA blood kit with a proteinase K digestion at 60°C for an hour in lysis buffer.

### cfDNA extraction

cfDNA was extracted using the Apostle MiniMax High Efficiency cfDNA Extraction kit (Apostle Bio, San Jose, CA). Briefly, extractions were performed in a 24-well deep well plate with the entire volume of patient plasma (average: 8 mL) in aliquots of up to 3mL. For aliquot volumes of less than a full milliliter, PBS, pH 7.4 was used to supplement volumes up to the next full milliliter. Total cfDNA was quantified using the Qubit fluorometer, and select samples were checked using Agilent TapeStation D1000 High Sensitivity or cfDNA assay.

### Duplex Library Preparation

Duplex libraries were prepared from matched normal gDNA, tumor gDNA, and cfDNA. For gDNA, 20-30ng gDNA was fragmented using the NEBNext FS module. The fragmentation reaction was processed directly into end repair, and end repair for all samples was performed using the KAPA HyperPrep kit (Roche, Indianapolis, IN). cfDNA inputs were normalized to 20 or 30ng. When the cfDNA concentration was too low to achieve 30ng, the maximum volume (50uL) of cfDNA was used for library preparation. Ligation was performed using the KAPA reagents with a pool of custom duplex adaptors (IDT). Each adaptor contained a UMI of 5 or 6 non-random nucleotides, and the pool contained adaptors with a Hamming distance of at least two. After ligation, the samples were purified with SPRI beads (Beckman Coulter, Indianapolis, IN) prior to indexing. Indexing was performed with unique dual indices (IDT) before a final cleanup. Shotgun libraries were quantified using a Qubit fluorometer.

### Hybrid Capture and Sequencing

Patient-specific hybrid capture panels were designed to capture all coding variants present in the archival biopsy used for neoantigen prediction. For capture efficiency and footprint consistency, pools of multiple patients were designed together such that each pool contained probes specific to variants from six to nine patients. Therefore, each patient sample was captured using both its own set of probes and multiple other patient probes.

A static, universal panel was designed to capture common driver variants and positions of approximately 250 common SNPs for fingerprinting. The universal panel also captures the entire coding sequences of *TP53, PTEN, ARID1A, JAK1, JAK2, IFNGR1, IFNGR2, B2M, NLRC5, TAP1, TAP2*, and the HLA canonical class I genes: *HLA-A, HLA-B*, and *HLA-C*. The patient and universal panels were designed and manufactured using custom design for xGen Lockdown probes from IDT.

Shotgun libraries were captured overnight with both the corresponding patient-specific panel and the universal panel or exome baits (IDT). Following the manufacturer’s instructions, probes were incubated overnight with shotgun libraries, Human Cot DNA, and blockers at 65°C, and bead washes were performed the next day using IDT xGen hybrid capture reagents. Captured libraries were amplified using KAPA HiFi HotStart ReadyMix. Final libraries were quantified using Qubit, normalized, and pooled. Libraries were sequenced on an Illumina NovaSeq 6000 SP, S1, S2, or S4 kit using 2×151 (v1 reagents) or 2×157 (v1.5 reagents) read lengths.

### Bioinformatics Processing

FASTQ files from WES of tumor tissue and blood samples were aligned to hg38 with Burrows-Wheeler Aligner (BWA-MEM, version 0.7.13), duplicate alignments were marked with Picard (version 2.7.1), base quality score recalibration was performed with GATK (version 3.5), and variant calling was done by FreeBayes (version 1.0.2). HLA typing was performed with OptiType (version 1.2). FASTQ files from RNA-Seq of tumor-tissue were aligned with STAR (version 2.5.1b) and gene expression quantification was performed with RSEM (version 1.2.31) Variant calls, HLA typing and gene expression values were used for neoantigen prediction using the EDGE™ model as previously described (22). The top 20 predictions are included in each patient’s individualized vaccine.

For analysis of cfDNA sequencing data, the duplex UMI was trimmed from the FASTQ file of the paired reads and appended to the read names. Trimmed reads were aligned to hg38 using BWA-MEM (version 0.7.17). To create consensus reads, fgbio (https://github.com/fulcrumgenomics/fgbio, version 1.1.0) was used to group the primary reads based on each read’s duplex UMI and primary BAM alignments. Consensus reads were created requiring a duplex depth of at least 3x for each strand. Consensus reads were mapped back to hg38 using BWA-MEM. Sequencing metrics and target coverage were computed using Picard (version 2.7.1). Variant calling for cfDNA targeted monitoring was performed using FreeBayes (version 1.3.1) and VarDict (VarDictJava, version 1.6.0) followed by manual curation. For analytical characterization of the ctDNA monitoring assay, variant calling on remnant patient samples was performed using VarDict. Manual curation was performed after variant calling for removal of artifacts and errors.

### HLA Loss of Heterozygosity

HLA analysis was performed by first extracting reads aligning to the class I HLA region on chromosome 6, the *TAP1* and *TAP2* regions, and unaligned reads after initial mapping to hg38. A patient-specific reference for the HLA alleles was created using alleles reported by OptiType from gDNA extracted from whole blood and confirmed by third-party HLA typing (UCLA Immunogenetics Center). The reads were re-aligned to the patient-specific reference for *HLA-A, HLA-B*, and *HLA-C* using BWA-MEM. *TAP1* and *TAP2* were added to evaluate variants and private SNPs. The original duplex UMI information was transferred to the re-aligned reads using GATK, and fgbio was used to group and create consensus reads from the duplex UMI and patient-specific alignment information. Consensus reads were mapped back to a patient-specific reference.

HLA allele balance was calculated using the read count attributed to each patient allele as calculated by Picard. Each allele ratio was compared to the allele ratios in the patient’s gDNA using a Chi-square test for each HLA gene. Variant calling was performed using VarDict and manually curated.

### Assay Characterization

For orthogonal assessment, remnant cfDNA underwent whole exome sequencing. Twenty to fifty nanograms of matched genomic DNA and cfDNA samples were used to prepare libraries. High molecular weight genomic DNA samples were enzymatically fragmented for 20 minutes and then end repaired by using the NEBNext Ultra II FS module. cfDNA samples were end repaired by NEBNext Ultra II End Repair/dA-Tailing module. Ligation was performed using NEBNext II Ligation module with NEBNext Adaptor for Illumina on both genomic DNA and cfDNA samples and then digested with NEB USER enzyme for 15 minutes. A cleanup was performed on the post-ligated and digested DNA. Samples were indexed using IDT’s unique dual indices and NEBNext Ultra II Q5 Master Mix. A cleanup was performed on the post-indexed samples. Libraries were quantified using Qubit Fluorometer and Qubit 1X dsDNA High-Sensitivity assay. For each library preparation, a non-template control (NTC) was begun at the earliest step (enzymatic fragmentation or end repair). For all libraries, 2000ng was used for hybrid capture enrichment using IDT xGen hybrid capture reagents. Briefly, Cot-1 DNA and TruSeq blockers were combined with each library and lyophilized under heat and vacuum. Libraries were hybridized overnight at 65°C with hybridization buffer, hybridization enhancer, and IDT exome v2 biotinylated probes. The following day, libraries were captured by incubation with magnetic streptavidin-conjugated beads for at least one hour. The captured libraries were washed and amplified per IDTs manufacturer’s protocol. Final post-captured libraries were quantified using Qubit Fluorometer and Qubit 1X dsDNA High-Sensitivity assay (Thermo Fisher Scientific). Exome-enriched libraries were normalized, pooled, and sequenced on a NovaSeq 6000 using 2×151 read length.

Twenty-five dilutions were prepared from eleven remnant patient cfDNA samples diluted into cfDNA extracted from healthy donors. Two sets of patient-specific baits were designed to cover the variants in eight and seven patients, including two variant sets for patients not used in the characterization. Variants from two patients were used in both sets of patient baits from two independent designs to assess performance across different designs. Six healthy donors were sequenced with their matched gDNA. Whole blood was procured through a prospective collection from Stage IV CRC or NSCLC patients (BioIVT, Westbury, NY), irrespective of variant or treatment status, for additional negative controls unrelated to the patient designs. Libraries were prepared as described in *Duplex Library Preparation* with three operators and sequenced across S1, S2, and S4 flow cells on a NovaSeq 6000.

The variants from WES were compared for accuracy and linearity was examined across the VAF range of the individual variants. The mean variant allele frequency for each sample was determined using the VAF of all patient-specific variants targeted by the assay, regardless of detection in either method. To understand the dynamic range of the mean VAF measurement measured by the assay, linearity was also established across the range of sample VAFs that were tested. Linear regressions were fit using Graphpad Prism 9. The logit fit for limit of detection was performed using the python package statsmodels.

For the input study, three replicates were prepared from 2.5, 5, 10, 20, 30, and 50ng cfDNA for two remnant dilutions, and triplicate gDNA libraries were prepared with 30ng gDNA. Libraries were enriched with the appropriate patient panel and sequenced using two kits on a NovaSeq 6000. Single libraries of healthy donor cfDNA were prepared with 2.5, 5, 10, 20, and 30ng. All other libraries were prepared with 30-50ng cfDNA and 30ng gDNA. gDNA from buffy coat collected from the healthy donors was sequenced to subtract the background of the healthy donors used for dilution.

Accuracy was assessed by comparing the variants found through whole exome sequencing compared to the cfDNA assay. Sensitivity and specificity were calculated using the true positive and true negative variants in whole exome sequencing compared to the cfDNA monitoring assay. Intra-run libraries included the 20, 30, and 50ng replicates of the input study, and two additional remnant dilutions were used to prepare libraries for reproducibility. Inter-run libraries were prepared from three remnant dilutions on separate days with at least two operators. Two of the three were also assessed with different panel sets performed on different days. Precision was assessed by comparing the concordant variant calls of the targeted variants across the multiple replicates.

## References

1. Bronkhorst AJ, Ungerer V, Holdenrieder S. The emerging role of cell-free DNA as a molecular marker for cancer management. Biomol Detect Quantif 2019;17:100087 doi 10.1016/j.bdq.2019.100087.

2. Parikh AR, Van Seventer EE, Siravegna G, Hartwig AV, Jaimovich A, He Y, et al. Minimal Residual Disease Detection using a Plasma-only Circulating Tumor DNA Assay in Patients with Colorectal Cancer. Clinical Cancer Research 2021;27(20):5586–94 doi 10.1158/1078-0432.Ccr-21-0410.

3. Tie J, Cohen JD, Lahouel K, Lo SN, Wang Y, Kosmider S, et al. Circulating Tumor DNA Analysis Guiding Adjuvant Therapy in Stage II Colon Cancer. N Engl J Med 2022;386(24):2261–72 doi 10.1056/NEJMoa2200075.

4. Anagnostou V, Forde PM, White JR, Niknafs N, Hruban C, Naidoo J, et al. Dynamics of Tumor and Immune Responses during Immune Checkpoint Blockade in Non-Small Cell Lung Cancer. Cancer Res 2019;79(6):1214–25 doi 10.1158/0008-5472.CAN-18-1127.

5. Bratman SV, Yang SYC, Iafolla MAJ, Liu Z, Hansen AR, Bedard PL, et al. Personalized circulating tumor DNA analysis as a predictive biomarker in solid tumor patients treated with pembrolizumab. Nature Cancer 2020;1(9):873–81 doi 10.1038/s43018-020-0096-5.

6. Goldberg SB, Narayan A, Kole AJ, Decker RH, Teysir J, Carriero NJ, et al. Early Assessment of Lung Cancer Immunotherapy Response via Circulating Tumor DNA. Clinical Cancer Research 2018;24(8):1872–80 doi 10.1158/1078-0432.Ccr-17-1341.

7. Hsu H-C, Lapke N, Wang C-W, Lin P-Y, You JF, Yeh CY, et al. Targeted Sequencing of Circulating Tumor DNA to Monitor Genetic Variants and Therapeutic Response in Metastatic Colorectal Cancer. Molecular Cancer Therapeutics 2018;17(10):2238–47 doi 10.1158/1535-7163.Mct-17-1306.

8. Thompson JC, Carpenter EL, Silva BA, Rosenstein J, Chien AL, Quinn K, et al. Serial Monitoring of Circulating Tumor DNA by Next-Generation Gene Sequencing as a Biomarker of Response and Survival in Patients With Advanced NSCLC Receiving Pembrolizumab-Based Therapy. JCO Precision Oncology 2021(5):510–24 doi 10.1200/po.20.00321.

9. Pellini B, Madison RW, Childress MA, Miller ST, Gjoerup O, Cheng J, et al. Circulating Tumor DNA Monitoring on Chemo-immunotherapy for Risk Stratification in Advanced Non–Small Cell Lung Cancer. Clinical Cancer Research 2023;29(22):4596–605 doi 10.1158/1078-0432.CCR-23-1578.

10. Vega DM, Nishimura KK, Zariffa N, Thompson JC, Hoering A, Cilento V, et al. Changes in Circulating Tumor DNA Reflect Clinical Benefit Across Multiple Studies of Patients With Non-Small-Cell Lung Cancer Treated With Immune Checkpoint Inhibitors. JCO Precis Oncol 2022;6:e2100372 doi 10.1200/PO.21.00372.

11. Zhang Q, Luo J, Wu S, Si H, Gao C, Xu W, et al. Prognostic and Predictive Impact of Circulating Tumor DNA in Patients with Advanced Cancers Treated with Immune Checkpoint Blockade. Cancer Discovery 2020;10(12):1842–53 doi 10.1158/2159-8290.Cd-20-0047.

12. Parikh AR, Leshchiner I, Elagina L, Goyal L, Levovitz C, Siravegna G, et al. Liquid versus tissue biopsy for detecting acquired resistance and tumor heterogeneity in gastrointestinal cancers. Nat Med 2019;25(9):1415–21 doi 10.1038/s41591-019-0561-9.

13. Finkle JD, Boulos H, Driessen TM, Lo C, Blidner RA, Hafez A, et al. Validation of a liquid biopsy assay with molecular and clinical profiling of circulating tumor DNA. npj Precision Oncology 2021;5(1):1–12 doi doi:10.1038/s41698-021-00202-2.

14. Rose Brannon A, Jayakumaran G, Diosdado M, Patel J, Razumova A, Hu Y, et al. Enhanced specificity of clinical high-sensitivity tumor mutation profiling in cell-free DNA via paired normal sequencing using MSK-ACCESS. Nature Communications 2021;12(1):1–12 doi doi:10.1038/s41467-021-24109-5.

15. Woodhouse R, Li M, Hughes J, Delfosse D, Skoletsky J, Ma P, et al. Clinical and analytical validation of FoundationOne Liquid CDx, a novel 324-Gene cfDNA-based comprehensive genomic profiling assay for cancers of solid tumor origin. PLoS One 2020;15(9):e0237802 doi 10.1371/journal.pone.0237802.

16. Lanman RB, Mortimer SA, Zill OA, Sebisanovic D, Lopez R, Blau S, et al. Analytical and Clinical Validation of a Digital Sequencing Panel for Quantitative, Highly Accurate Evaluation of Cell-Free Circulating Tumor DNA. PLOS ONE 2015;10(10):e0140712 doi 10.1371/journal.pone.0140712.

17. Jia Q, Chiu L, Wu S, Bai J, Peng L, Zheng L, et al. Tracking Neoantigens by Personalized Circulating Tumor DNA Sequencing during Checkpoint Blockade Immunotherapy in Non-Small Cell Lung Cancer. Adv Sci (Weinh) 2020;7(9):1903410 doi 10.1002/advs.201903410.

18. Abbosh C, Birkbak NJ, Wilson GA, Jamal-Hanjani M, Constantin T, Salari R, et al. Phylogenetic ctDNA analysis depicts early-stage lung cancer evolution. Nature 2017;545(7655):446-51 doi 10.1038/nature22364.

19. Davis AA, Iams WT, Chan D, Oh MS, Lentz RW, Peterman N, et al. Early Assessment of Molecular Progression and Response by Whole-genome Circulating Tumor DNA in Advanced Solid Tumors. Molecular Cancer Therapeutics 2020;19(7):1486–96 doi 10.1158/1535-7163.Mct-19-1060.

20. Zviran A, Schulman RC, Shah M, Hill STK, Deochand S, Khamnei CC, et al. Genome-wide cell-free DNA mutational integration enables ultra-sensitive cancer monitoring. Nature Medicine 2020;26(7):1114–24 doi doi:10.1038/s41591-020-0915-3.

21. Wong D, Luo P, Oldfield L, Gong H, Brunga L, Rabinowicz R, et al. Integrated analysis of cell-free DNA for the early detection of cancer in people with Li-Fraumeni Syndrome. medRxiv 2022:2022.10.07.22280848 doi 10.1101/2022.10.07.22280848.

22. Bulik-Sullivan B, Busby J, Palmer CD, Davis MJ, Murphy T, Clark A, et al. Deep learning using tumor HLA peptide mass spectrometry datasets improves neoantigen identification. Nature Biotechnology 2018;37(1):55–63 doi doi:10.1038/nbt.4313.

23. Palmer CD, Rappaport AR, Davis MJ, Hart MG, Scallan CD, Hong S-J, et al. Individualized, heterologous chimpanzee adenovirus and self-amplifying mRNA neoantigen vaccine for advanced metastatic solid tumors: phase 1 trial interim results. Nature Medicine 2022;28(8):1619–29 doi doi:10.1038/s41591-022-01937-6.

24. Loupakis F, Sharma S, Derouazi M, Murgioni S, Biason P, Rizzato MD, et al. Detection of Molecular Residual Disease Using Personalized Circulating Tumor DNA Assay in Patients With Colorectal Cancer Undergoing Resection of Metastases. JCO Precision Oncology 2021(5):1166–77 doi 10.1200/PO.21.00101.

25. Reinert T, Henriksen TV, Christensen E, Sharma S, Salari R, Sethi H, et al. Analysis of Plasma Cell-Free DNA by Ultradeep Sequencing in Patients With Stages I to III Colorectal Cancer. JAMA Oncol 2019;5(8):1124–31 doi 10.1001/jamaoncol.2019.0528.

26. Zhao J, Reuther J, Scozzaro K, Hawley M, Metzger E, Emery M, et al. Personalized Cancer Monitoring Assay for the Detection of ctDNA in Patients with Solid Tumors. Molecular Diagnosis & Therapy 2023:1–16 doi doi:10.1007/s40291-023-00670-1.

27. Abbosh C, Frankell AM, Harrison T, Kisistok J, Garnett A, Johnson L, et al. Tracking early lung cancer metastatic dissemination in TRACERx using ctDNA. Nature 2023;616(7957):553-62 doi doi:10.1038/s41586-023-05776-4.

28. Al Bakir M, Huebner A, Martínez-Ruiz C, Grigoriadis K, Watkins TBK, Pich O, et al. The evolution of non-small cell lung cancer metastases in TRACERx. Nature 2023;616(7957):534-42 doi doi:10.1038/s41586-023-05729-x.

29. Kennedy SR, Schmitt MW, Fox EJ, Kohrn BF, Salk JJ, Ahn EH, et al. Detecting ultralow-frequency mutations by Duplex Sequencing. Nat Protoc 2014;9(11):2586–606 doi 10.1038/nprot.2014.170.

30. Anderson P, Aptsiauri N, Ruiz-Cabello F, Garrido F. HLA class I loss in colorectal cancer: implications for immune escape and immunotherapy. Cell Mol Immunol 2021;18(3):556–65 doi 10.1038/s41423-021-00634-7.

31. D’Amico S, Tempora P, Melaiu O, Lucarini V, Cifaldi L, Locatelli F, et al. Targeting the antigen processing and presentation pathway to overcome resistance to immune checkpoint therapy. Frontiers in Immunology 2022;13 doi 10.3389/fimmu.2022.948297.

32. Dhatchinamoorthy K, Colbert JD, Rock KL. Cancer Immune Evasion Through Loss of MHC Class I Antigen Presentation. Frontiers in Immunology 2021;12 doi 10.3389/fimmu.2021.636568.

33. Gettinger S, Choi J, Hastings K, Truini A, Datar I, Sowell R, et al. Impaired HLA Class I Antigen Processing and Presentation as a Mechanism of Acquired Resistance to Immune Checkpoint Inhibitors in Lung Cancer. Cancer Discovery 2017;7(12):1420–35 doi 10.1158/2159-8290.Cd-17-0593.

34. Sade-Feldman M, Jiao YJ, Chen JH, Rooney MS, Barzily-Rokni M, Eliane J-P, et al. Resistance to checkpoint blockade therapy through inactivation of antigen presentation. Nature Communications 2017;8(1):1–11 doi doi:10.1038/s41467-017-01062-w.

35. Abbosh C, Swanton C, Birkbak NJ. Clonal haematopoiesis: a source of biological noise in cell-free DNA analyses. Ann Oncol 2019;30(3):358–9 doi 10.1093/annonc/mdy552.

36. Jaiswal S, Ebert BL. Clonal hematopoiesis in human aging and disease. Science 2019;366(6465) doi 10.1126/science.aan4673.

37. Salk JJ, Loubet-Senear K, Maritschnegg E, Valentine CC, Williams LN, Higgins JE, et al. Ultra-Sensitive TP53 Sequencing for Cancer Detection Reveals Progressive Clonal Selection in Normal Tissue over a Century of Human Lifespan. Cell Rep 2019;28(1):132–44 e3 doi 10.1016/j.celrep.2019.05.109.

38. JS B, PA W, P C. Pathways of antigen processing. Annual review of immunology 2013;31 doi 10.1146/annurev-immunol-032712-095910.

39. A K, J T. Genetics of antigen processing and presentation. Immunogenetics 2019;71(3) doi 10.1007/s00251-018-1082-2.

40. Ritz U, Seliger B. The Transporter Associated With Antigen Processing (TAP): Structural Integrity, Expression, Function, and Its Clinical Relevance. Molecular Medicine 2001;7(3):149–58 doi doi:10.1007/BF03401948.

41. Hastings RK, Openshaw MR, Vazquez M, Moreno-Cardenas AB, Fernandez-Garcia D, Martinson L, et al. Longitudinal whole-exome sequencing of cell-free DNA for tracking the co-evolutionary tumor and immune evasion dynamics: longitudinal data from a single patient. Ann Oncol 2021;32(5):681–4 doi 10.1016/j.annonc.2021.02.007.

42. McGranahan N, Rosenthal R, Hiley CT, Rowan AJ, Watkins TBK, Wilson GA, et al. Allele-Specific HLA Loss and Immune Escape in Lung Cancer Evolution. Cell 2017;171(6):1259–71 e11 doi 10.1016/j.cell.2017.10.001.

43. Marijt KA, Griffioen L, Blijleven L, van der Burg SH, van Hall T. Cross-presentation of a TAP-independent signal peptide induces CD8 T immunity to escaped cancers but necessitates anchor replacement. Cancer Immunology, Immunotherapy 2021;71(2):289–300 doi doi:10.1007/s00262-021-02984-7.

44. Oliveira CC, van Hall T. Alternative Antigen Processing for MHC Class I: Multiple Roads Lead to Rome. Frontiers in Immunology 2015;6 doi 10.3389/fimmu.2015.00298.

45. Keenan TE, Burke KP, Van Allen EM. Genomic correlates of response to immune checkpoint blockade. Nature Medicine 2019;25(3):389–402 doi 10.1038/s41591-019-0382-x.

46. Sucker A, Zhao F, Pieper N, Heeke C, Maltaner R, Stadtler N, et al. Acquired IFNγ resistance impairs anti-tumor immunity and gives rise to T-cell-resistant melanoma lesions. Nat Commun 2017;8:15440 doi 10.1038/ncomms15440.

47. Zaretsky JM, Garcia-Diaz A, Shin DS, Escuin-Ordinas H, Hugo W, Hu-Lieskovan S, et al. Mutations Associated with Acquired Resistance to PD-1 Blockade in Melanoma. https://doiorg/101056/NEJMoa1604958 2016 doi NJ201609013750906.

48. FDA U. Use of Circulating Tumor Deoxyribonucleic Acid for Early-Stage Solid Tumor Drug Development; Draft Guidance for Industry; Availability | FDA. US FDA; 2022.

49. Anagnostou V, Ho C, Nicholas G, Juergens RA, Sacher A, Fung AS, et al. ctDNA response after pembrolizumab in non-small cell lung cancer: phase 2 adaptive trial results. Nat Med 2023;29(10):2559–69 doi 10.1038/s41591-023-02598-9.

50. Lheureux S, Prokopec SD, Oldfield LE, Gonzalez-Ochoa E, Bruce JP, Wong D, et al. Identifying Mechanisms of Resistance by Circulating Tumor DNA in EVOLVE, a Phase II Trial of Cediranib Plus Olaparib for Ovarian Cancer at Time of PARP Inhibitor Progression. Clinical Cancer Research 2023:OF1–OF11 doi 10.1158/1078-0432.Ccr-23-0797.

