## Supplemental Figures for "Comprehensive Longitudinal ctDNA Monitoring in Metastatic Cancer Patients Treated with an Individualized Neoantigen-directed Vaccine"

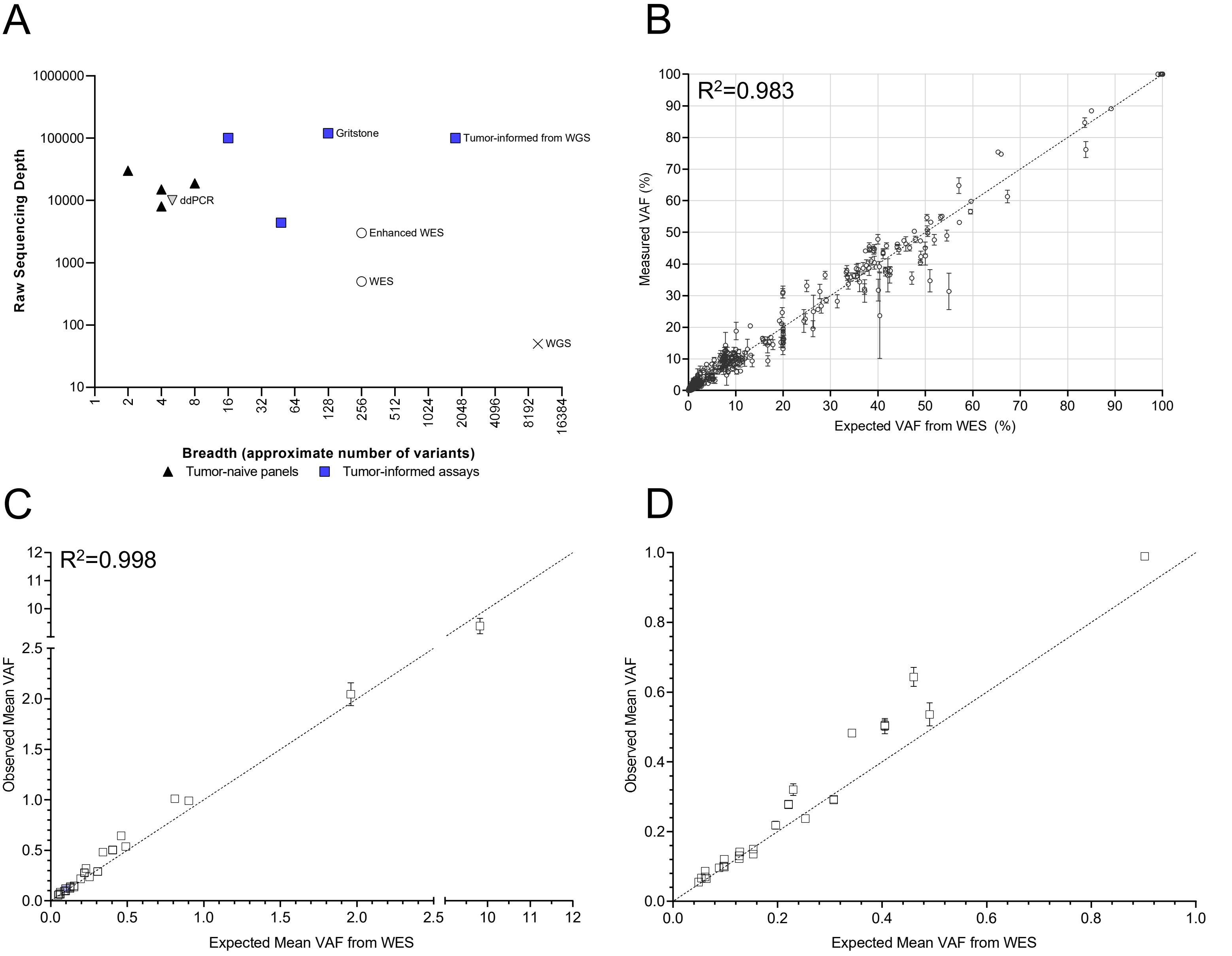


Supplemental Figure 1. A) ctDNA monitoring assays vary both their depth and breadth. Tumor-naïve panels target the smallest number of variants. Tumor-informed variants can result in increased specificity with the addition of more targeted regions. B) Variant-level linearity of the ctDNA monitoring assay. Including somatic and germline variants from 0.02-100% VAF, the ctDNA assay variant detection and linearity was evaluated against whole exome sequencing. Line of identity is shown, and symbols and error bars represent mean ± standard deviation. C) Linearity of cfDNA quantification using mean VAF. Remnant samples were diluted into healthy donors ranging from 0.049-9.82% and mean VAF was calculated as measured using the ctDNA assay. Line of identity is shown, and symbols and error bars represent mean ± standard deviation. D) Linearity of ctDNA quantification in samples with mean VAF ≤1% (lower left portion of Supplementary Figure 1C). Remnant clinical cfDNA samples were diluted into healthy donor cfDNA. The measured VAF of the ctDNA assay was compared to mean VAF by WES. Line of identity is shown, and symbols and error bars represent mean ± standard deviation.

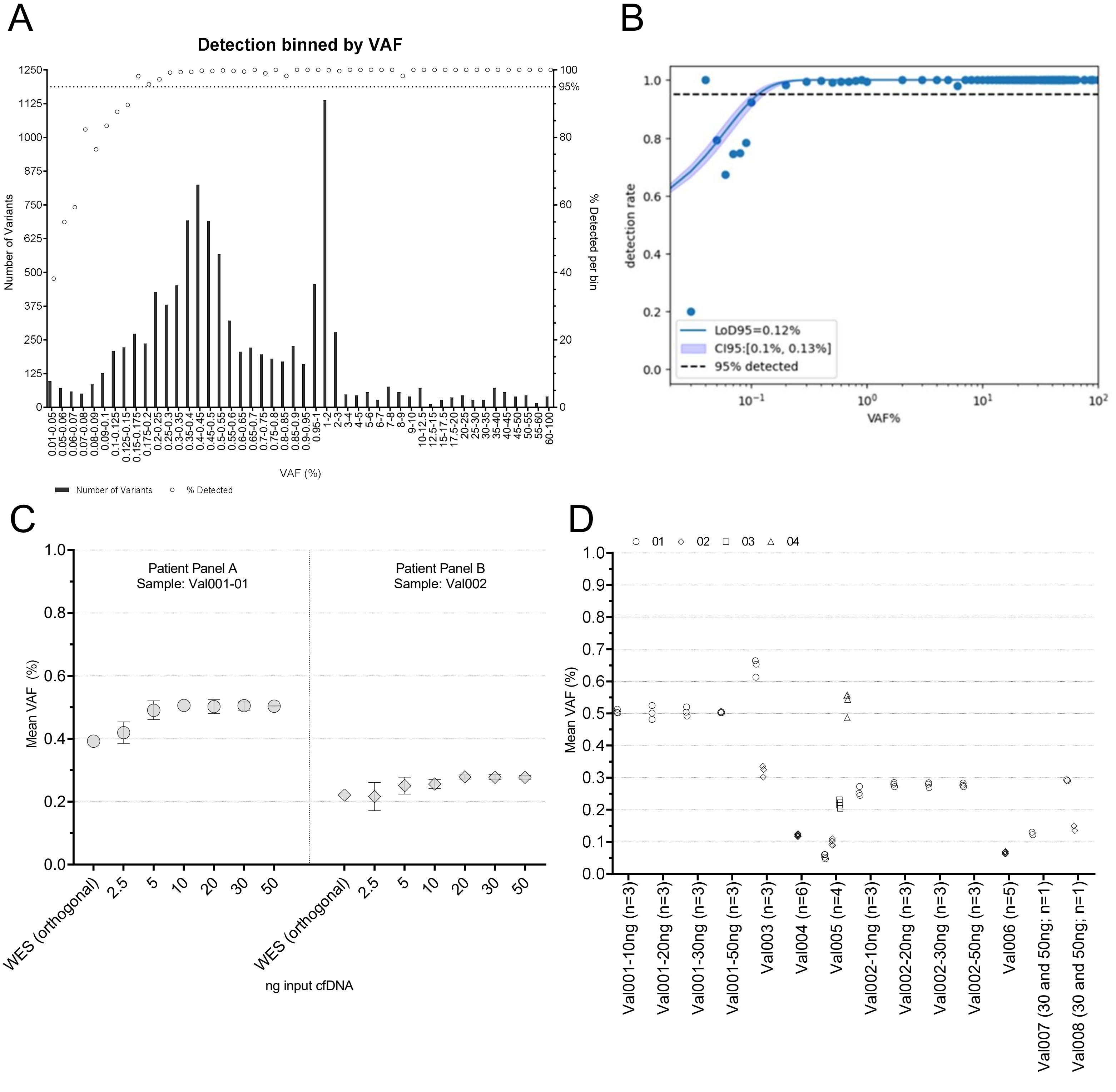


Supplemental Figure 2. A) Variants were categorized by their VAF, and the detection frequency of each bin was calculated. B) A Logit regression is fit to the detection rate as a function of VAF% at a per-variant level across the replicates. The 95% detection level occurred at 0.12% VAF for a single variant. C) To determine the precision across cfDNA inputs, two samples (Val0001-01 and Val002) were prepared with three replicates from 2.5-50ng. Val0001-01 was captured using patient panel A, and Val002 was captured using patient panel B. Mean VAF was consistent across inputs and among replicates, varying the greatest at the lowest cfDNA input. Symbols and error bars represent mean ± standard deviation. D) Replicate samples were prepared across operators, sequencing runs, reagent lots, and patient panel designs, producing consistent measurements of mean VAF.

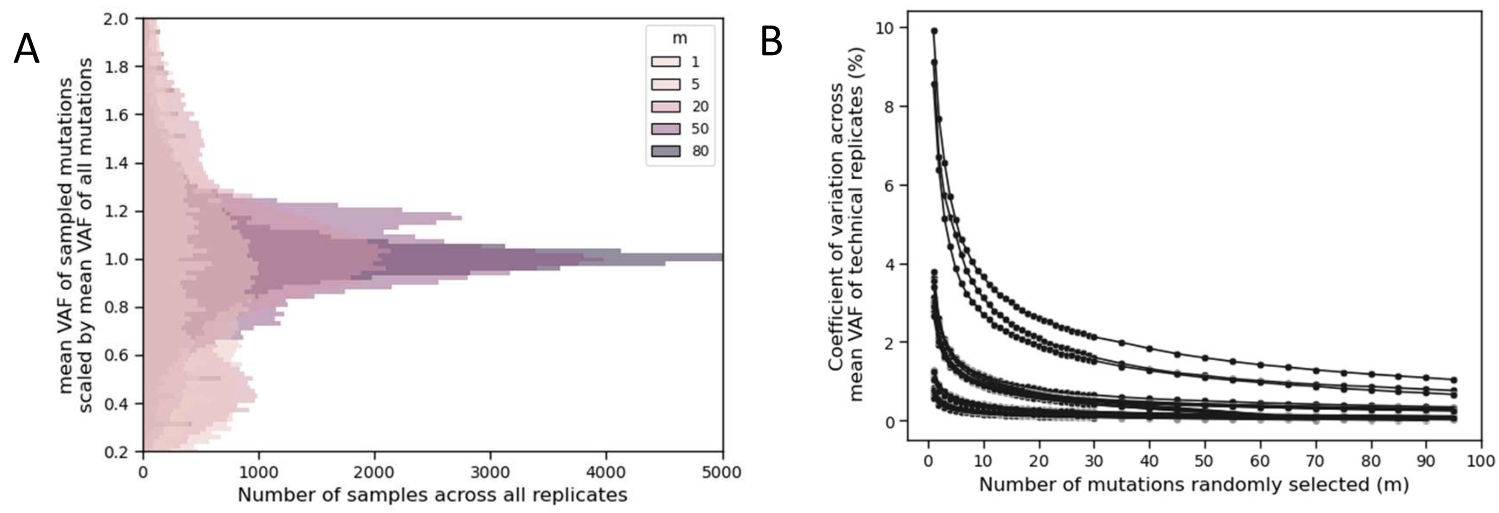


Supplemental Figure 3. Mean VAF variability based on number of variants used in the mean VAF calculation. The validation technical replicates were used in this analysis. For every number of mutations (m), each replicate was sampled 1000 times. A) Visualization of the mean VAF for the random samples with a specific number of mutations (m) compared to all mutations. As a larger fraction of the variants are selected, the mean VAF converges to the expected mean VAF of the targeted variants (behaving similarly to Chebyshev's inequality). B) The coefficient of variation (CV) for all samples, calculated across the replicates. The CV decreases as more variants are selected. Each line represents a set of technical replicates from the remnant dilutions.

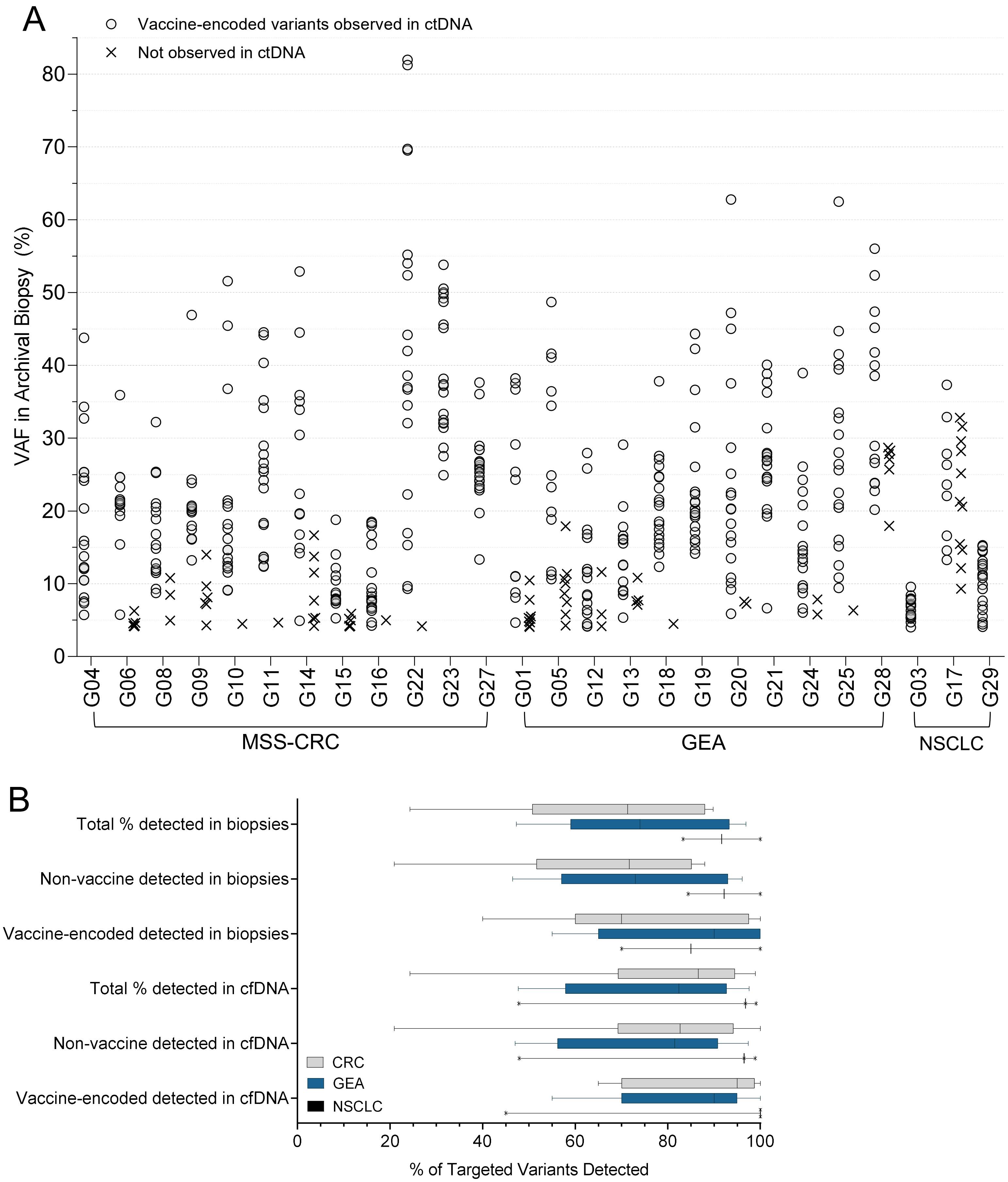


Supplemental Figure 4. A) Distribution of VAF of vaccine-encoded variants in the archival biopsy and their detection in ctDNA. B) The percentage of vaccine-encoded variants and non-vaccine, targeted tumor variants were similar in both ctDNA and tissue biopsies across the three tissue types. G02 and G07 did not have biopsies collected due to no evidence of disease nor any variant detection in ctDNA.

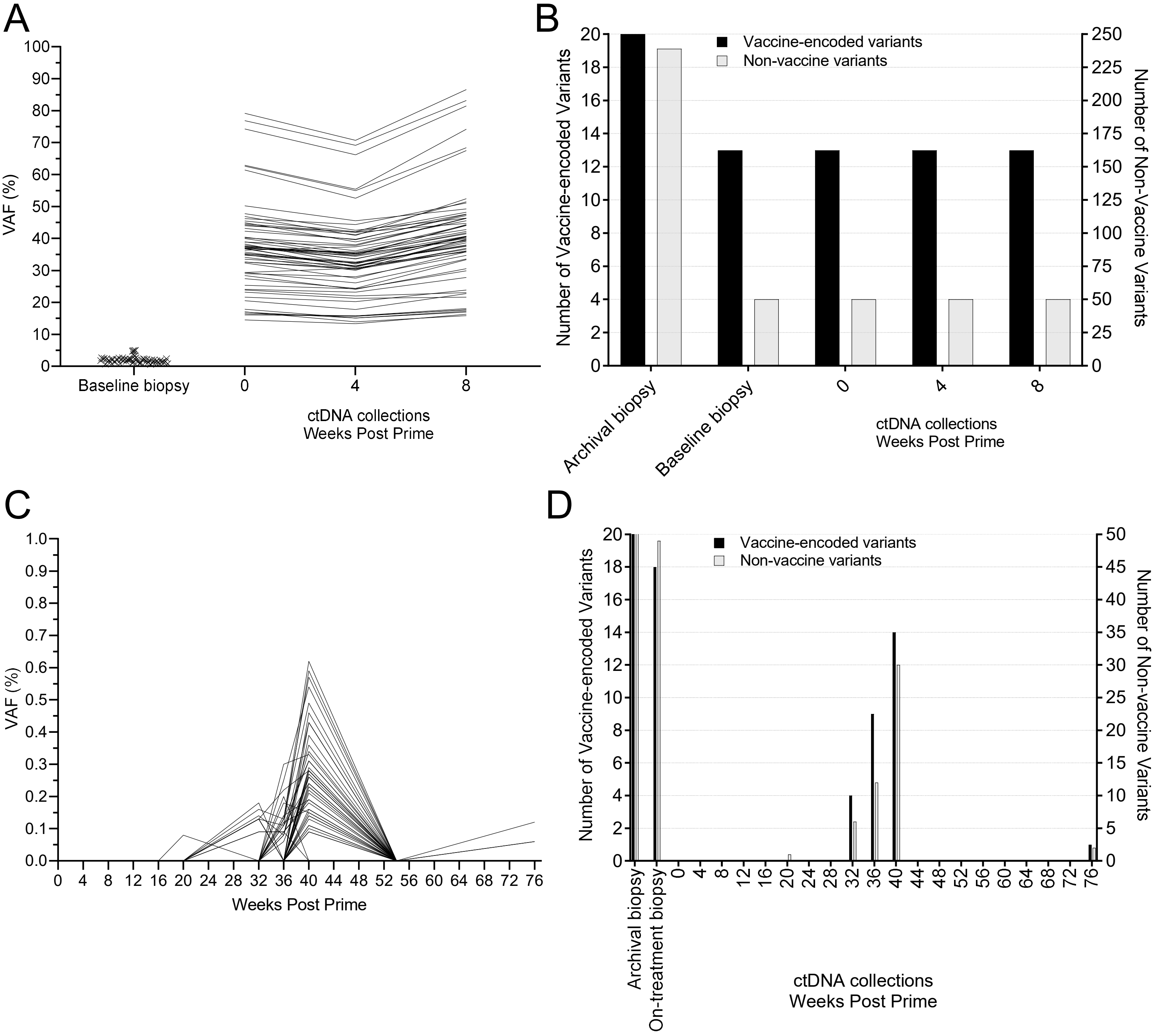


Supplemental Figure 5. A) Patient G15's biopsy contained low tumor content. ctDNA was detected for 63 variants that were used for longitudinal monitoring. B) G15 longitudinal variant monitoring showed an overlap of variants in both the baseline tissue biopsy and all ctDNA samples that were collected. The majority of the nearly 250 targeted variants in the archival biopsy were not detected in either the subsequent tissue or ctDNA. C) Patient G13 ctDNA dynamics showed ctDNA negativity for the first 16 weeks before becoming positive at 20 weeks. The VAF peaked at 40 weeks and subsequently dropped, but the patient continued to be ctDNA positive. D) Patient G13 was one of two patients where the variant detection in ctDNA was less than that of the available tissue biopsies. The number of variants detected in the tissue biopsy was consistently higher than in ctDNA.

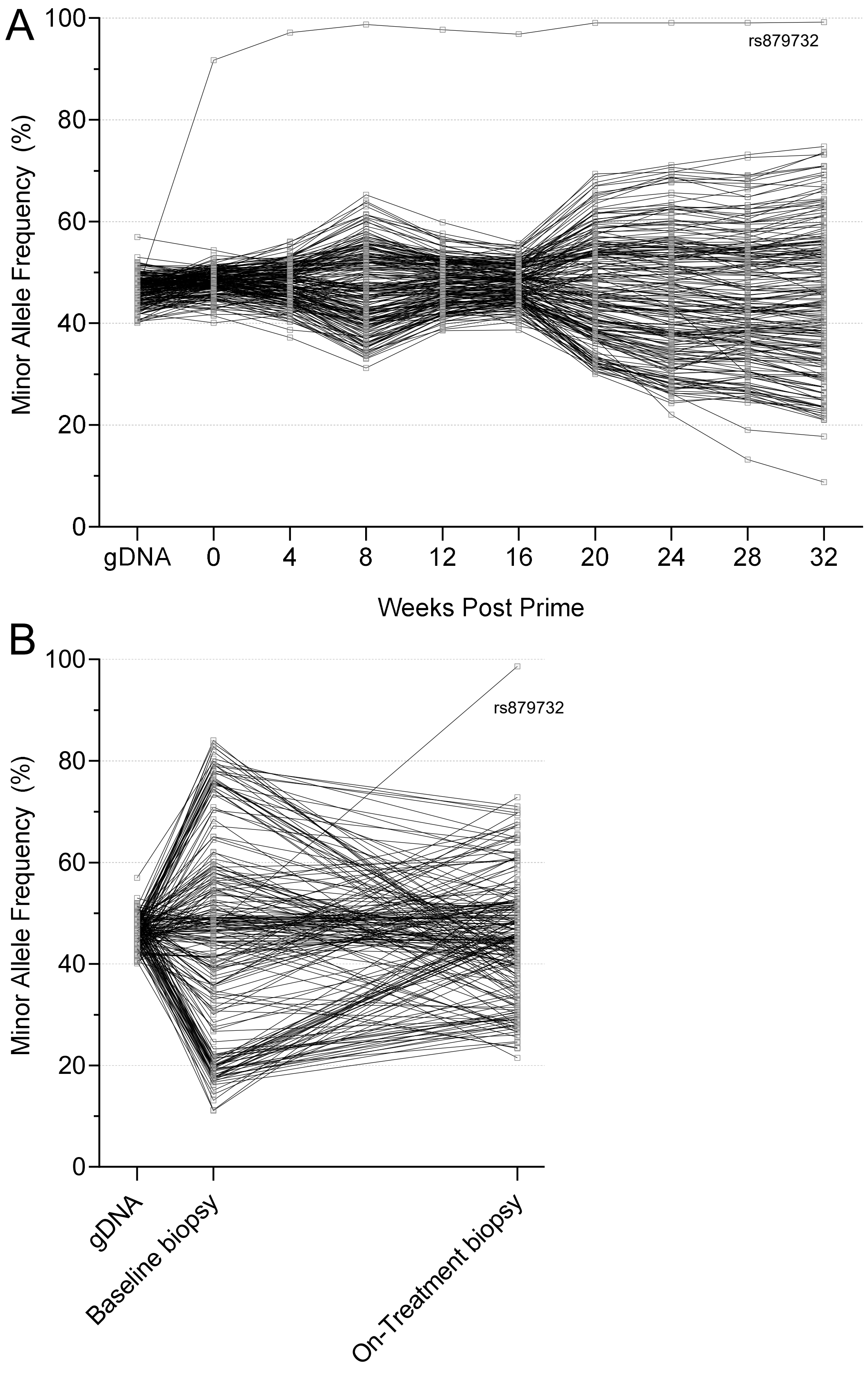


Supplemental Figure 6. Longitudinal monitoring of B-allele frequency (BAF) of heterozygous SNPs in patient G24. A) BAF analysis of cfDNA demonstrated an increasing imbalance and genomic instability. The position of SNP rs879732 achieved near homozygosity in ctDNA 4 weeks after starting treatment. B) Similar to the cfDNA, the baseline biopsy and on-treatment biopsy of patient G24 had genomic instability and changes in allelic balance at heterozygous positions. The position for SNP rs879732 did not show the same LOH in the baseline biopsy (47.2% VAF) that was seen in the cfDNA (>90% VAF) and on-treatment biopsy (98.7%).

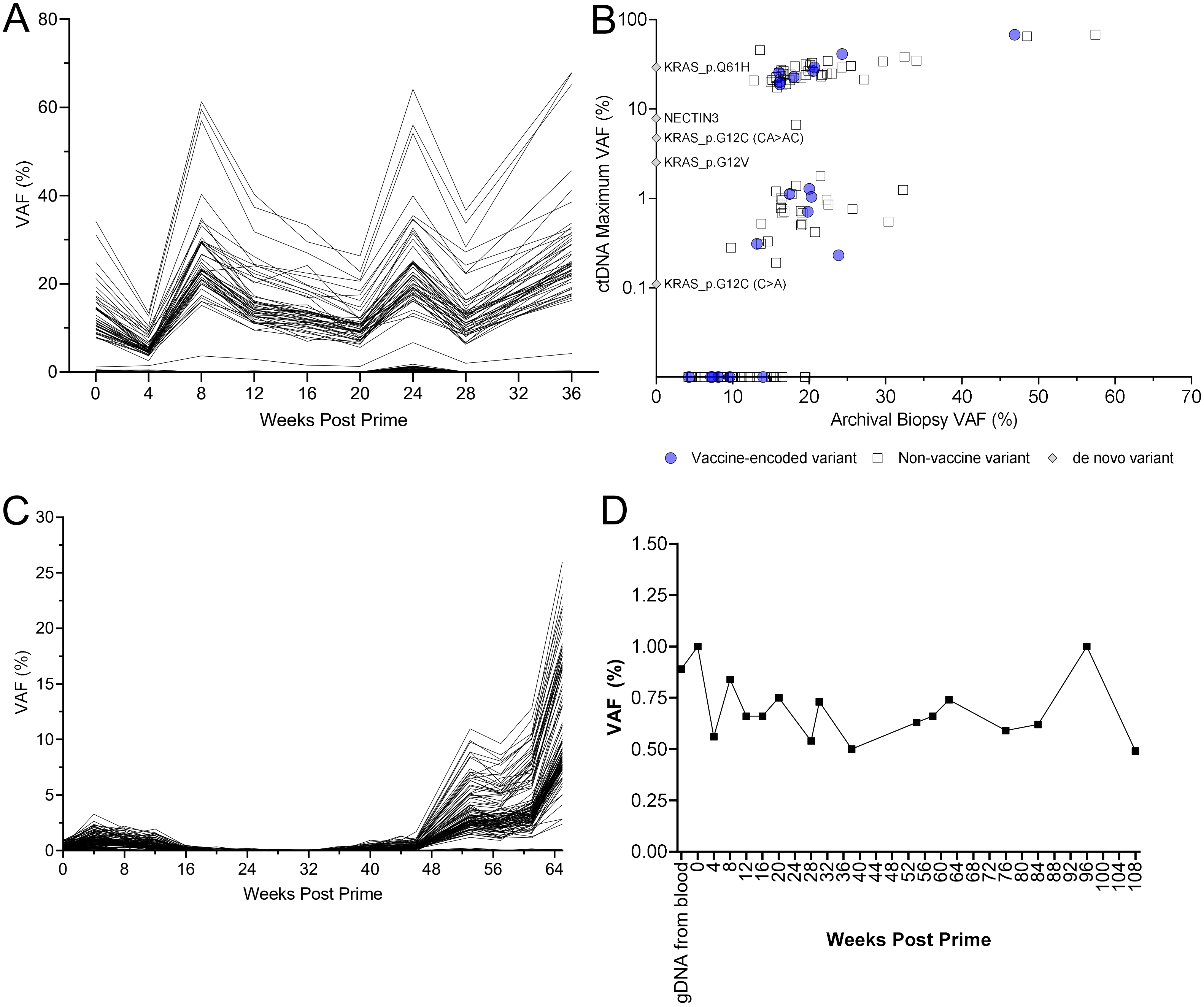


Supplemental Figure 7. A) G09 ctDNA dynamics of targeted patient-specific variants. B) Comparison of VAF in ctDNA and archival biopsy in patient G09 indicates distinct populations of variants, including *de novo* variants. Filled blue dots indicate cassette variants. Gray diamonds are newly-detected variants. C) G08 ctDNA dynamics of targeted, patient-specific variants showed clearance of ctDNA before increasing slowly at 40 weeks. D) Detection of clonal hematopoiesis from sequencing of gDNA from blood. CHIP variants, a source of biological noise, can confound somatic variant calling.

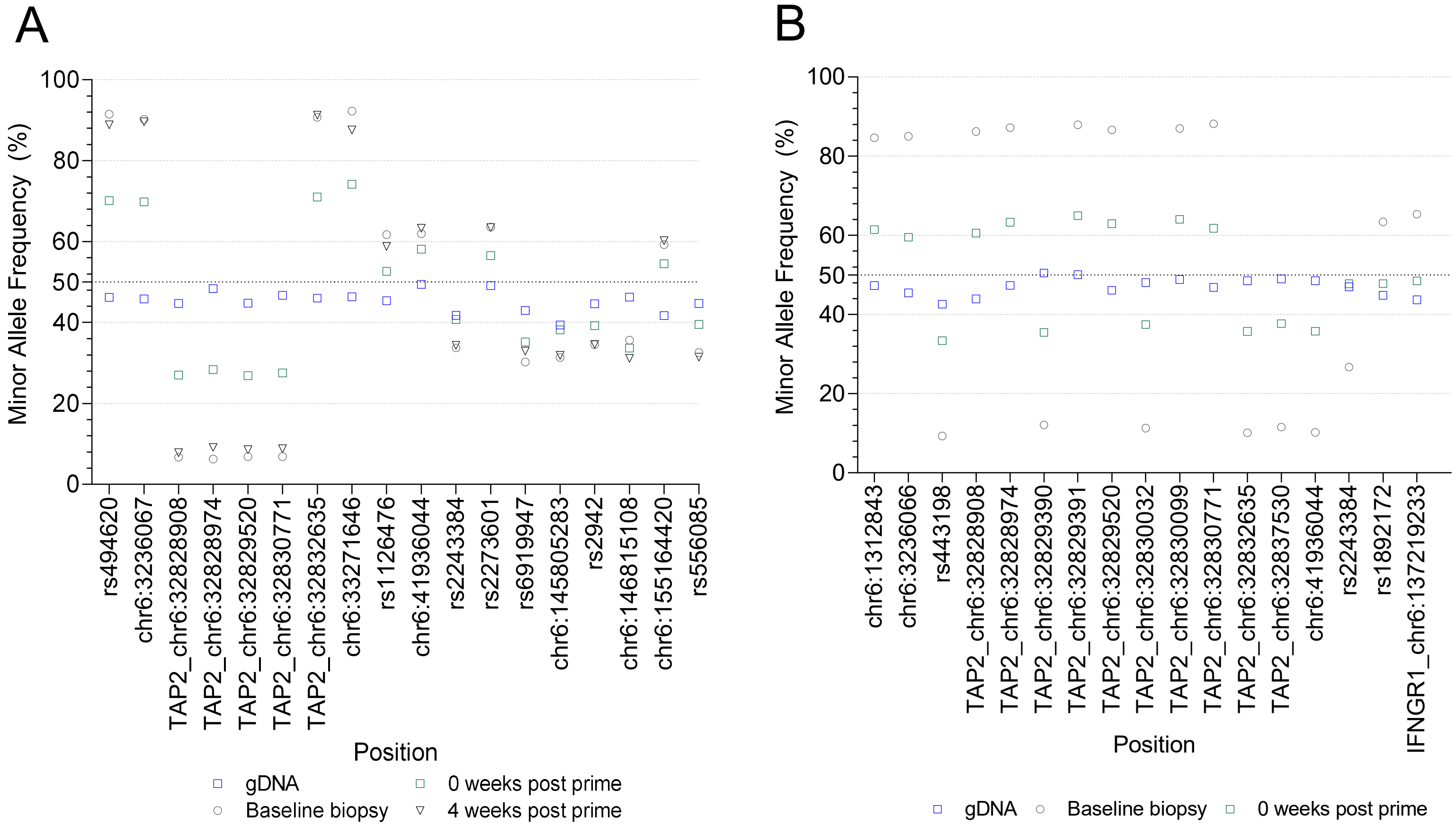


Supplemental Figure 8. Two MSS-CRC patients contained evidence of HLA haplotype loss in both tissue biopsies and cfDNA. B-allele frequency along chr6 showed expanded LOH beyond the HLA locus in both patients. A) Patient G04 B) Patient G23

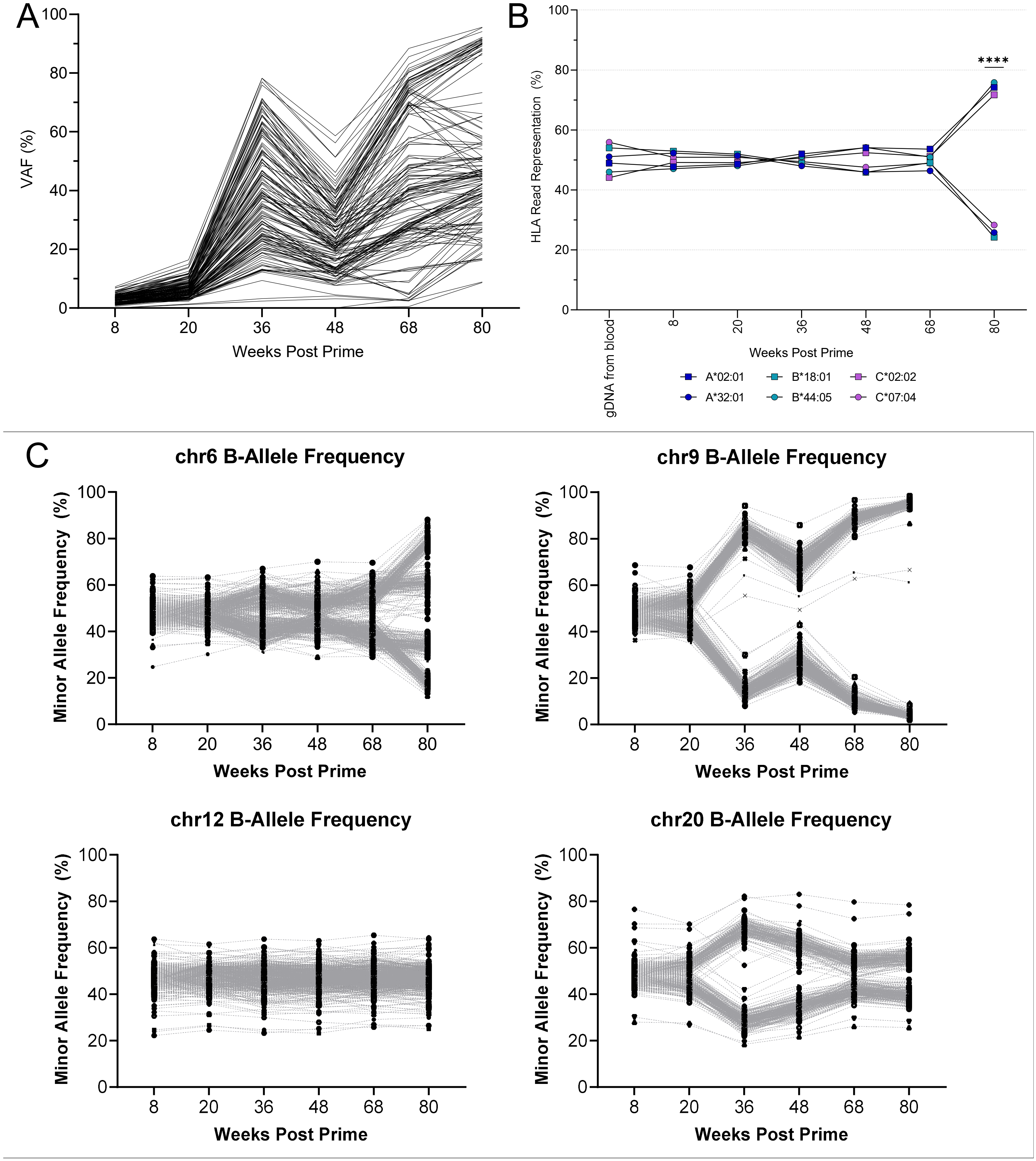


Supplemental Figure 9. Whole exome sequencing of select ctDNA samples from G01. A) The targeted archival variants from WES of G01 ctDNA mirrored the dynamics observed from the ctDNA assay B) Evidence of HLA LOH from WES of cfDNA samples was present at 80 weeks. C) LOH at positions of heterozygous SNPs of select chromosomes, including chromosome 6, the location of HLA class I, showed various levels of LOH across the genome.

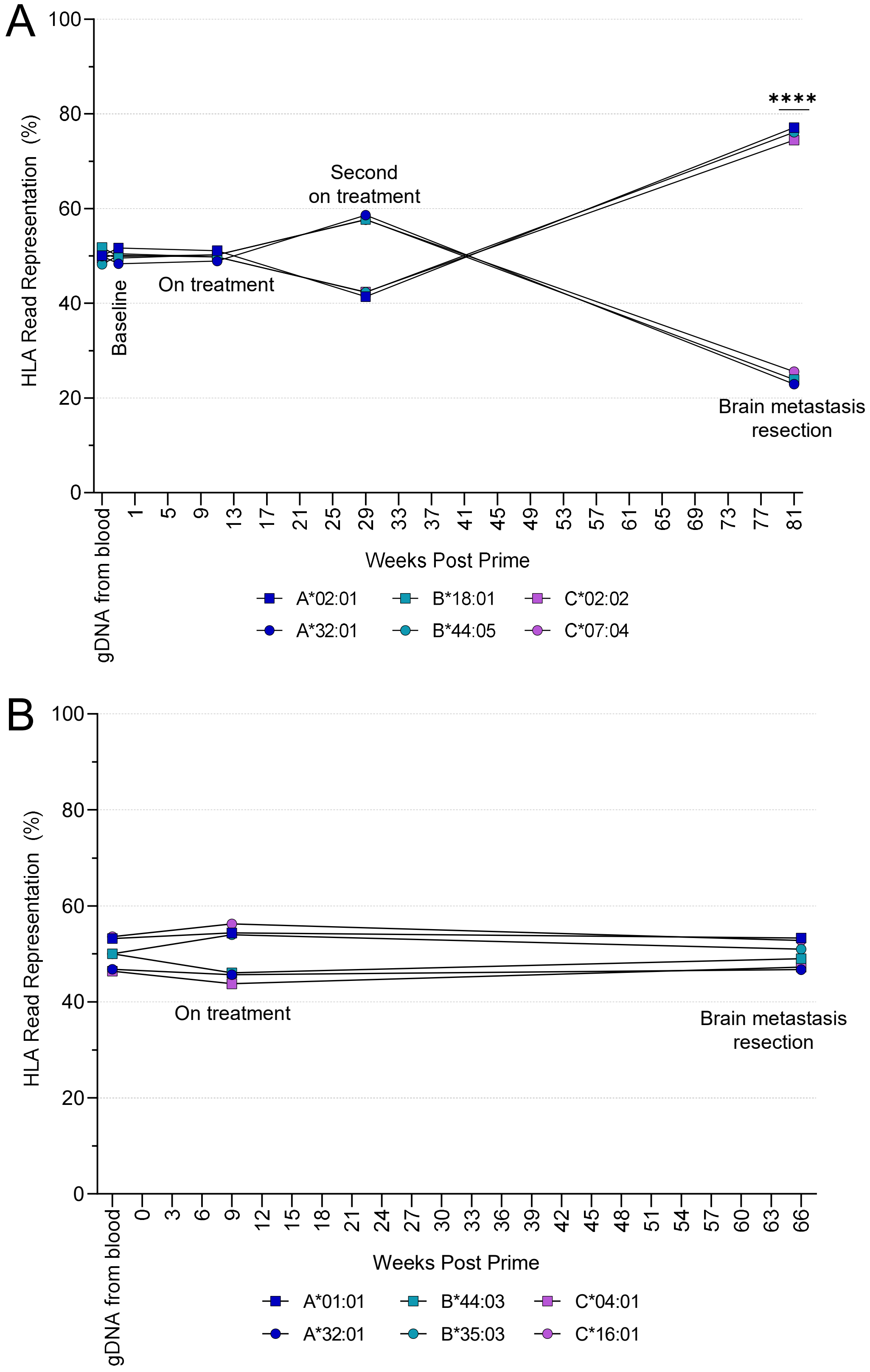


Figure S10. A) WES of the tissue from the brain metastasis of G01 contained HLA LOH that was also found in the patient's cfDNA. B) G16 FFPE biopsies did not have evidence of HLA LOH as observed in the cfDNA.
